# A unified analysis of cell-type and trajectory-associated pathways in single-cell data using Phoenix

**DOI:** 10.1101/2025.09.29.679361

**Authors:** Yudit Halperin, Daphna Nachmani, Michal Rabani

## Abstract

Single-cell RNA sequencing has transformed our ability to resolve complex cellular heterogeneity within biospecimens at the molecular level. However, identifying which biological pathways accurately reflect distinct cell types or continuous cellular trajectories remains a major challenge. Traditional methods often miss subtle or non-linear pathway activities, limiting biological interpretability and insights. To address this, we develop *Phoenix*, a pathway analysis framework that leverages random forest models and non-parametric significance testing to evaluate the relevance of functional gene sets for distinguishing between cell-types and organizing cells along pseudotemporal cellular trajectories. *Phoenix* reveals both up- and downregulated processes, including those shaped by complex non-linear gene interactions, and quantifies their effect sizes. Applied to human and mouse hematopoiesis as well as zebrafish embryogenesis, *Phoenix* identifies both cell-type-specific and trajectory-associated pathways, spanning housekeeping, developmental, and lineage-specific programs. It outperforms existing tools in capturing cell-type-specific activities of small pathways and reveals greater overlap in pathway activities across species. Ultimately, *Phoenix* provides a sensitive and interpretable framework for uncovering biologically meaningful pathways and eliciting the interactions between their components in complex single-cell datasets, opening new opportunities to explore dynamic gene regulation across biological systems.

## Introduction

Advances in single-cell RNA sequencing (scRNA-seq) technologies have enabled high-resolution molecular profiling of heterogeneous biological samples at unprecedented scale (Papalexi and Satija 2018; Farrell et al. 2018; Tusi et al. 2018; Ranzoni et al. 2021; Tirosh and Suva 2024; Potter 2018). As datasets grow in size and complexity, extracting meaningful biological insights requires robust functional interpretation of gene expression data. Pathway analysis plays a central role in this effort, facilitating the interpretation of biological processes from gene-level signals in scRNA-seq.

Traditionally, functional interpretation has followed a “gene-centric” approach. These bottom-up methods first aggregate cells to identify “interesting” subsets of genes, such as differentially expressed genes with respect to an underlying structure (e.g., between cell types or conditions) and then assess their enrichment within predefined pathways (e.g., Gene Ontology (The Gene Ontology Consortium et al. 2000, 2023) or KEGG (Kanehisa et al. 2025; Kanehisa and Goto 2000)). Gene set enrichment analysis (GSEA) and related tools (Mootha et al. 2003; Subramanian et al. 2005), originally developed for bulk transcriptomic data, fall into this category. However, such methods are often challenged by the sparsity, noise, and high dimensionality inherent to single-cell data and are limited in their ability to capture the complex, dynamic, and heterogeneous nature of single-cell transcriptomes. Therefore, tools specifically developed for single-cell pathway enrichment aim to infer the activity of pathways within single-cells, by taking a more “pathway-centric” approach (Drier et al. 2013). Rather than relying on intermediate gene lists, these top-down methods directly infer activity scores of predefined, biologically relevant pathways at the resolution of individual cells. Tools such as AUCell (Aibar et al. 2017), scGSEA (Franchini et al. 2023) and ssGSEA (Barbie et al. 2009) preserve single-cell resolution, but remain susceptible to some sparsity and technical noise and are not always optimal. More recent methods, such as GSDensity (Liang et al. 2023) and SiPSic (Davis et al. 2024), have improved sensitivity for detecting pathway heterogeneity across cells. Nonetheless, they still rely on absolute changes in expression level and may overlook more subtle, but biologically meaningful, pathway dynamics as well as non-linear interactions between pathway components. Moreover, their cluster-independent outputs, resulting in per-cell activity levels, can be challenging to interpret without additional downstream analysis. Finally, tools such as TIPS (Zheng et al. 2021), have been introduced to analyze pathway activity along pseudotemporal trajectories (e.g., during differentiation or development). These methods are better suited for capturing the continuous nature of cellular state transitions, but do not address discrete cell-types.

Here, we introduce *Phoenix*, a computational framework designed to assess the relevance of functional pathways to both discrete cell identities and continuous pseudotemporal trajectories in single-cell data. *Phoenix* evaluates each pathway in the context of the intrinsic structure of the dataset, whether defined by discrete cell-types or continuous lineages, offering a unified and statistically robust approach to pathway relevance. *Phoenix* analysis is based on a supervised ensemble model (random forest), offering an interpretable approach that can recover non-linear gene interactions within pathways. Applied to diverse scRNA-seq datasets, including hematopoietic and developmental systems, *Phoenix* demonstrates improved sensitivity and interpretability over existing methods, establishing it as a powerful tool for uncovering biologically meaningful pathway activities in complex single-cell transcriptomic landscapes.

## Results

### *Phoenix*: a pathway analysis framework for scRNA-seq data

*Phoenix* (**Fig. 1**, **Methods**) assesses the relevance of functional pathways and other gene sets to both cell types and cellular trajectories in scRNA-seq experiments (**Fig. 1A**). It trains a random forest model on genes from a specific pathway, either for classification of single cells into cell types or for regression to predict pseudotime along a trajectory (**Fig. 1B**), facilitating a functional analysis of processes reflected both by discrete cell states and continuous progression. The resulting models also rank genes within the pathway by their relevance for the task. *Phoenix* evaluates pathway relevance with three complementary metrics. First, non-parametric significance testing compares each pathway’s performance to a background distribution of performance scores evaluated from randomly selected gene sets of a similar size (**Supplemental Fig. S1**). Pathways outperforming random gene sets in predicting cell types or pseudotime (FDR < 1%) highlight processes that distinguish cell types and organize cells along trajectories (**Fig. 1C**). When a large pathways set is used, background distributions can also be evaluated on real pathways (**Supplemental Fig. S2**). Second, an effect size metric quantifies the average pathway expression within versus outside a cell type (classification) or between earliest and later pseudotime cells (regression), also indicating whether pathways are up- or downregulated, and the pseudotime of maximal difference. Third, *Phoenix* analyzes per-gene contributions within the random-forest model to determine pathway-level behavior, by considering the fraction of pathway genes with low contributions.

**Figure 1.**
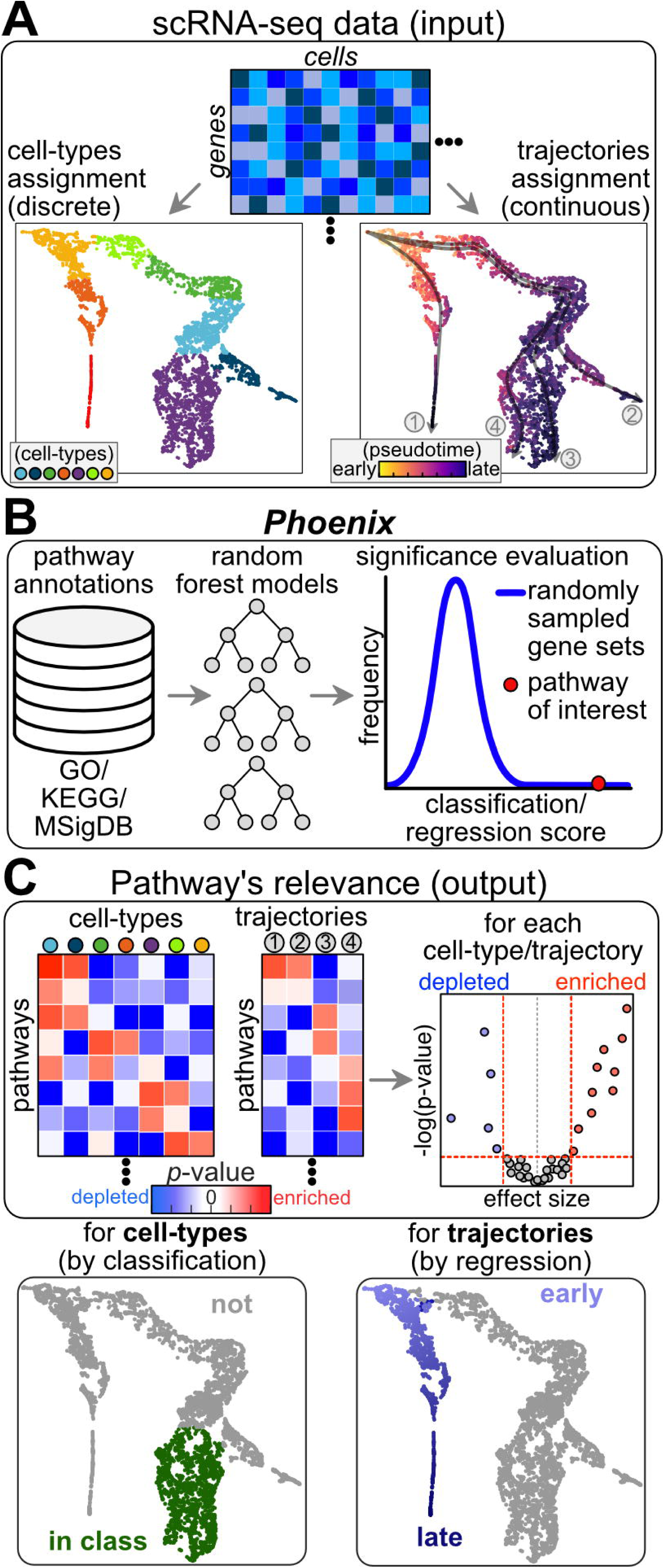
*Phoenix*: a pathway analysis framework for scRNA-seq data. *Phoenix* is a computational framework designed to assess the relevance of functional pathways to both discrete cell identities and continuous pseudotemporal trajectories in single-cell data. **(A)** *Phoenix* uses single-cell RNA-seq (scRNA-seq) data as input. A matrix of raw scRNA-seq counts (top) can be analyzed to either define discrete cell types (marked by colors, left) or to organize cells along developmental trajectories (marked by gray arrows 1-4) and predict pseudotime (marked by purple-yellow colorscale, right). Cell-types and trajectories can be either calculated by *Phoenix* from raw scRNA-seq counts, or provided as input (preferred). Each trajectory represents an inferred path of a dynamic biological process through the expression space, while pseudotime assigns each cell a relative position along that path, approximating progression without relying on real time measurements. One or more trajectories can exist within a dataset, depending on its composition. **(B)** *Phoenix* assesses the relevance of functional pathways and other gene lists (left) to both cell-types and cellular trajectories within a scRNA-seq experiment, by training a random forest model (middle) on genes from a specific pathway for classification of single cells into cell-types, or regression to predict cells’ pseudotime along a trajectory. The results are evaluated by a non-parametric significance testing, comparing the performance of each pathway to that of randomly selected sets of genes of the same size (right). **(C)** This analysis calculates a *p*-value and effect size for the ability of each pathway to distinguish each cell-type from other cells in the dataset (cell-type classification), or for the ability of each pathway to predict the organization of cells along a developmental trajectory (pseudotime regression). *Phoenix* reports the results of this analysis both as tables (left), and by volcano plots (one plot for each cell-type/trajectory, right) to reveal how effect sizes and *p*-values are balanced in the results. For each association, *Phoenix* further plots the combined expression of pathway genes, distinguishing associated cell-type (left) or reconstructing temporal progression along a trajectory (right).

*Phoenix* offers several important advantages for pathway analysis. First, it highlights pathways based on their ability to distinguish discrete classes (cell types) or predict continuous properties (pseudotimes) within a specific scRNA-seq experiment. This allows *Phoenix* to detect significant results even when changes are relatively subtle or when not all genes are equally informative, as long as they collectively provide strong separation between targets. To aid interpretation, *Phoenix* provides a ranked list of pathway genes based on per-gene contributions to the random-forest models learned, highlighting the key players involved in the relevant biological processes within a cell type. Contributions assigned to genes in significant pathways were higher, and had fewer low-importance genes (**Supplemental Fig. S3**). Second, by relying on a random forest model, *Phoenix* captures complex, non-linear interactions between genes in a pathway, making it well-suited for analyzing complex regulatory interactions. Random forest models also allow *Phoenix* to associate both positive and negative effects with cell types or trajectories, revealing gene sets that are either up- or downregulated within a specific cell population. Third, *Phoenix* leverages user-supplied trajectory and cell-type inference as target values for the analysis, which can be independently optimized for the dataset under study. This approach mitigates limitations of single-cell data, such as noisy estimation of expression for individual genes in individual cells, and provides enhanced statistical power to identify biologically relevant gene sets. Finally, *Phoenix* addresses both discrete and continuous aspects of single-cell data. This dual capability enables it to uncover pathways that are critical for distinguishing between cell types, as well as those involved in continuous processes such as cellular differentiation across pseudotime.

Consequently, *Phoenix* offers an interpretable framework for interrogating tissue physiology and heterogeneity with high sensitivity and accuracy. *Phoenix* is available as an open-source Python package, providing a well-documented easily installed tool for single-cell pathway analysis. Benchmarking analysis provides the user information on *Phoenix* runtime and memory requirements (**Supplemental Fig. S4**).

### *Phoenix* confirms existing knowledge of hematopoietic differentiation

We first validated *Phoenix* using scRNA-seq data of hematopoietic differentiation (**Supplemental Table S1**) from both human (Ranzoni et al. 2021) and mouse (Tusi et al. 2018). The human dataset comprises of 2,646 cells grouped into 12 clusters of progenitor and differentiated hematopoietic cell types based on marker analysis (**Fig. 2A**). Trajectory mapping with *Slingshot* (Street et al. 2018) identified 5 differentiation trajectories, which ordered progenitor and mature cell populations as expected (**Fig. 2A**). The mouse dataset includes 4,751 cells assigned to 8 hematopoietic cell-type clusters and 7 differentiation trajectories (**Fig. 2B**).

**Figure 2.**
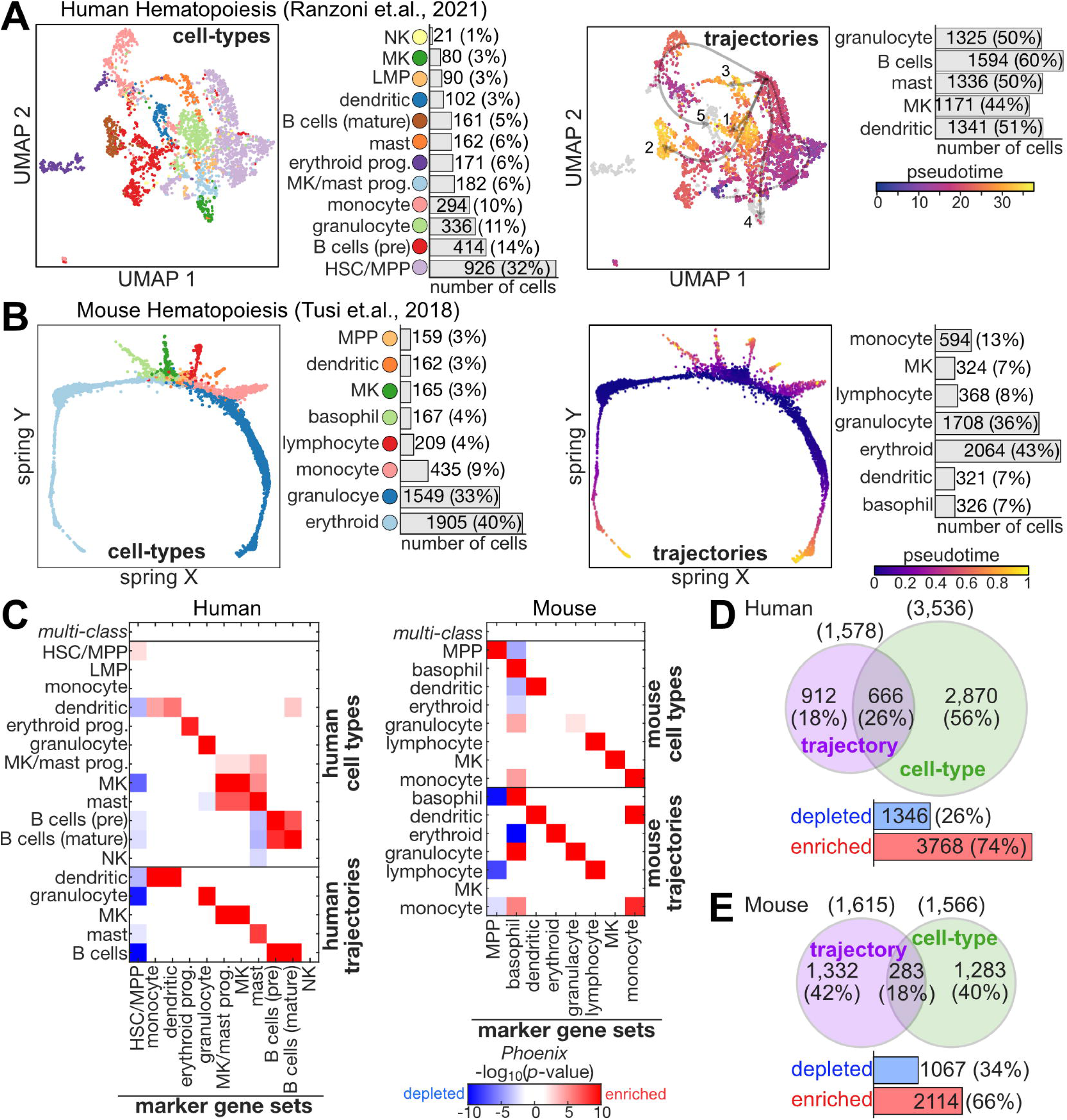
*Phoenix* confirms existing knowledge of hematopoietic differentiation. **(A-B)** Projection of human (Ranzoni et. al., 2021, UMAP, A) and mouse (Tusi et. al., 2018, Spring, B) hematopoiesis differentiation scRNA-seq data. Left: cells are colored by cell-type (as listed) and number of cells from each type is noted. Right: cells are colored by pseudotime (purple: early, yellow: late, gray: not assigned to a trajectory), trajectories are listed on right, with number of cells. For human trajectories in panel A, arrows indicate individual trajectories and are numbered according to their top-to-bottom order in the list on the right: trajectory 1 represents granulocytes, 2 B cells, 3 mast cells, 4 MK and 5 dendritic cells. The dotted line denotes the early segment that is shared among trajectories. **(C)** Matrix of *p*-values calculated by *Phoenix* for marker gene sets (columns) within cell-types (top) and trajectories (bottom) of hematopoietic scRNA-seq data (rows) reveals their association with expected cell-types within scRNA-seq data. First row of the matrix represents the classification of all cell-types in the data (multi-class classification). Color-scale represents significance (-log_10_(*p*-value)) and effect: red positive values for enriched pathways (positive effect size) and blue negative values for depleted pathways (negative effect size). Left: human, right: mouse data. **(D-E)** *Phoenix* enrichments in human (D) or mouse (E) datasets. Top: venn diagrams showing overlapping associations by *Phoenix* with cell-types or matching trajectories. Total number of associations and percentages are indicated. Bottom: counts of enriched (red, positive effect size) or depleted (blue, negative effect size) associations by *Phoenix*. Total number of associations and percentages are indicated.

Marker gene-sets (as defined in the original publications, **Supplemental Table S2**) enrichment confirmed our expectations. *Phoenix* identified highly significant associations for relevant marker set within the associated cell-types in each organism (**Fig. 2C**), as well as in a cross-comparison (**Supplemental Fig. S5A-B**). These results confirm the robustness of the observed associations, leaving only a minimal possibility of circularity between cell-type labels and the pathways tested.

Analysis of biological pathways in the MSigDB knowledgebase (Liberzon et al. 2011; Subramanian et al. 2005) (35,134 human, 16,276 mouse pathways) revealed a small fraction of significant associations, alleviating overfitting concerns. Because no external ground-truth pathway labels were available for benchmarking, we evaluated *Phoenix* performance using global statistics and expected biological relationships. In human data, *Phoenix* identified 5,115 (1%) significant associations (FDR < 1%, effect size > 1.2; **Supplemental Table S3**), primarily with cell types, and fewer with trajectories (**Fig. 2D**). In mouse data, *Phoenix* identified 3,181 (1.3%) significant associations (FDR < 1%, effect size > 1.2; **Supplemental Table S4**), with trajectories showing a similar number of associations as cell types (**Fig. 2E**). Enrichments were more common than depletions, although the balance varied across cell types and trajectories (**Fig. 2D-E, Supplemental Fig. S5C-D**). Number of significant associations also differed substantially between groups (cell types or trajectories) and was largely independent of cell-type size. However, large trajectories, such as the mouse granulocyte and erythroid lineages, exhibited particularly high numbers of significant associations (**Supplemental Fig. S5C-D**).

Human hematopoietic pathways reflected expected biology (**Fig. 3A**). Predictions associated lympho-myeloid progenitors (LMP) with spindle microtubules and cell-cycle programs; erythrocyte progenitors with oxygen transport and hemoglobin; megakaryocyte (MK) cells with alpha actinin binding (*Phoenix* volcano plots, **Fig. 3B**); mast cells and trajectory with acute inflammation and catechol/prostaglandin synthesis linked to runx1/runx1t targets (**Fig. 3C**); granulocytes with cytolytic granules and hydrogen peroxide metabolism; monocytes with chronic inflammation; dendritic cells with antigen processing and presentation (**Fig. 3D**); pre-B cells and the B cell trajectory with IgM immunoglobulin pathways (**Fig. 3E**); mature B cells with B cell differentiation.

**Figure 3.**
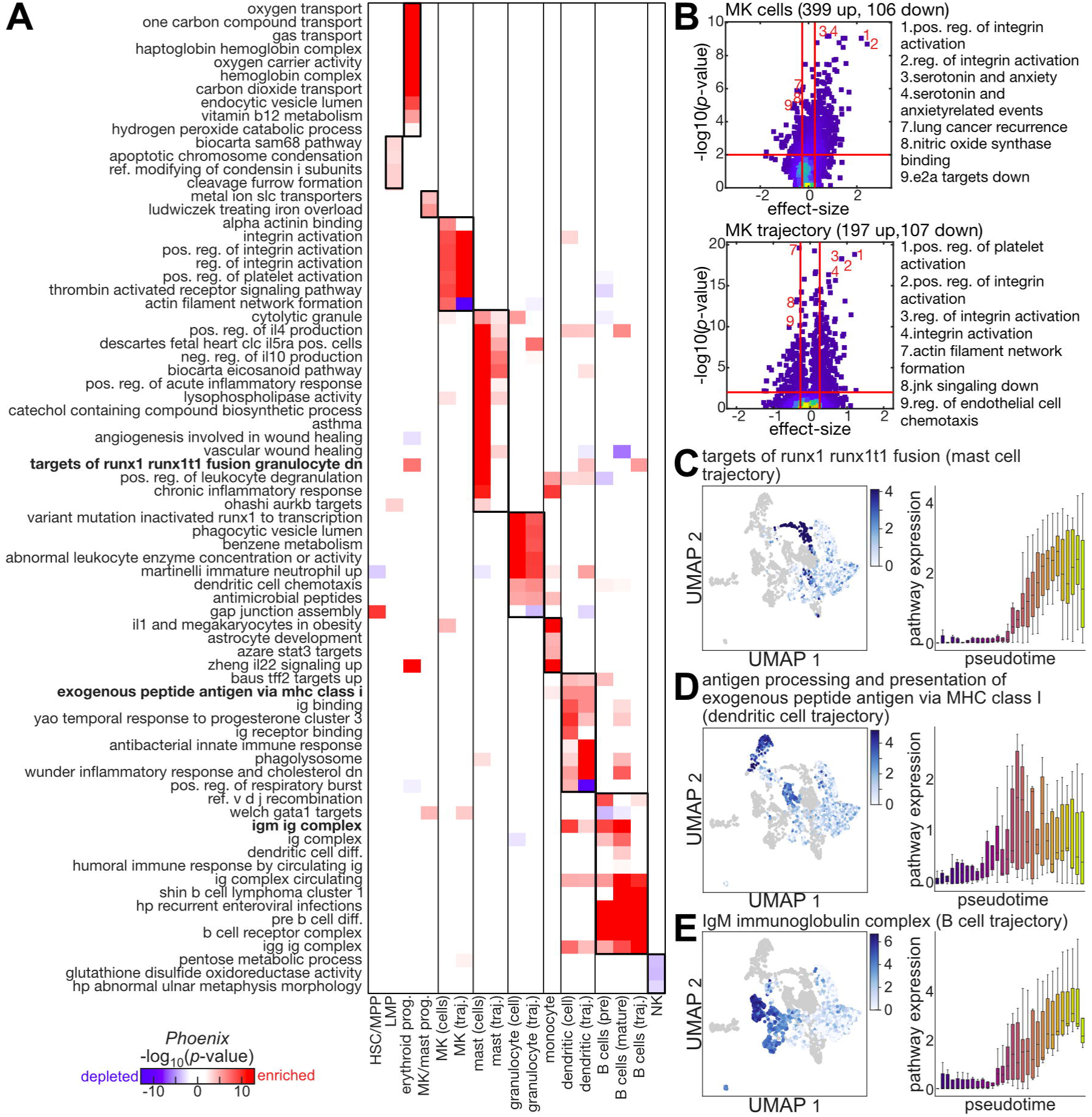
Pathways associated with human hematopoietic cell-types and trajectories. **(A)** Matrix of *p*-values calculated by *Phoenix* for the enrichment of top selected pathways (rows) within cell-types and trajectories of human hematopoietic scRNA-seq data (columns). Color-scale represents significance (-log_10_(*p*-value)) and effect: red positive values for enriched pathways (positive effect size) and blue negative values for depleted pathways (negative effect size). **(B)** Volcano plots showing *p*-values (y-axis, FDR-corrected, -log10 scale) and effect sizes (x-axis) of associations revealed by *Phoenix* in MK cells (left) and trajectory (right). Colors represent density (blue = low density, yellow = high density), and red lines represent significance thresholds (p<0.01, absolute effect > 1.2). Top selected positive and negative annotations are numbered and indicated next to plot. **(C-E)** Detailed expression data from selected pathways (as noted on top, and labeled in bold in panel (A)) that were associated with specific human hematopoietic differentiation trajectories by *Phoenix*. Top: scRNA-seq UMAP with trajectory cells colored by the average expression of genes within the selected pathway (blue color-scale, as indicated). Bottom: boxplot of average expression of genes within the selected pathway (y-axis) across pseudotime bins (x-axis). Box color represent pseudotime (purple: early; yellow: late). Central line represents the median, box edges are 25th and 75th percentiles, whiskers extend to largest/smallest value except outliers; outlier points are plotted individually.

Mouse hematopoietic data showed similar biologically relevant enrichments (**Fig. 4A**). *Phoenix* associated hematopoietic multipotent progenitor (MPP) cells with stem cell proliferation, and myeloid gene expression; erythroid trajectory with hemoglobin biosynthesis (**Fig. 4B**); MK cells with hedgehog signaling and depletion of heme biosynthesis; granulocyte trajectory with degranulation (positive) and acute inflammation (**Fig. 4C**); basophils with notch signaling and resposne to interferon; monocytes with neutrophil activation and inflammatory response, and azurophil granules; dendritic cells with exogenous antigen processing and presentation (**Fig. 4D**), and dendritic cell differentiation; lymphocyte trajectory with interleukin production; lymphocyte cells with DNA modification (**Fig. 4E**) and DNA-directed DNA polymerase activity. Mouse enrichments also revealed noticeable differences in which pathways were associated with cell-types or trajectories of the same cell population (**Fig. 4A**), reflecting possible distinctions in the underlying biological process.

**Figure 4.**
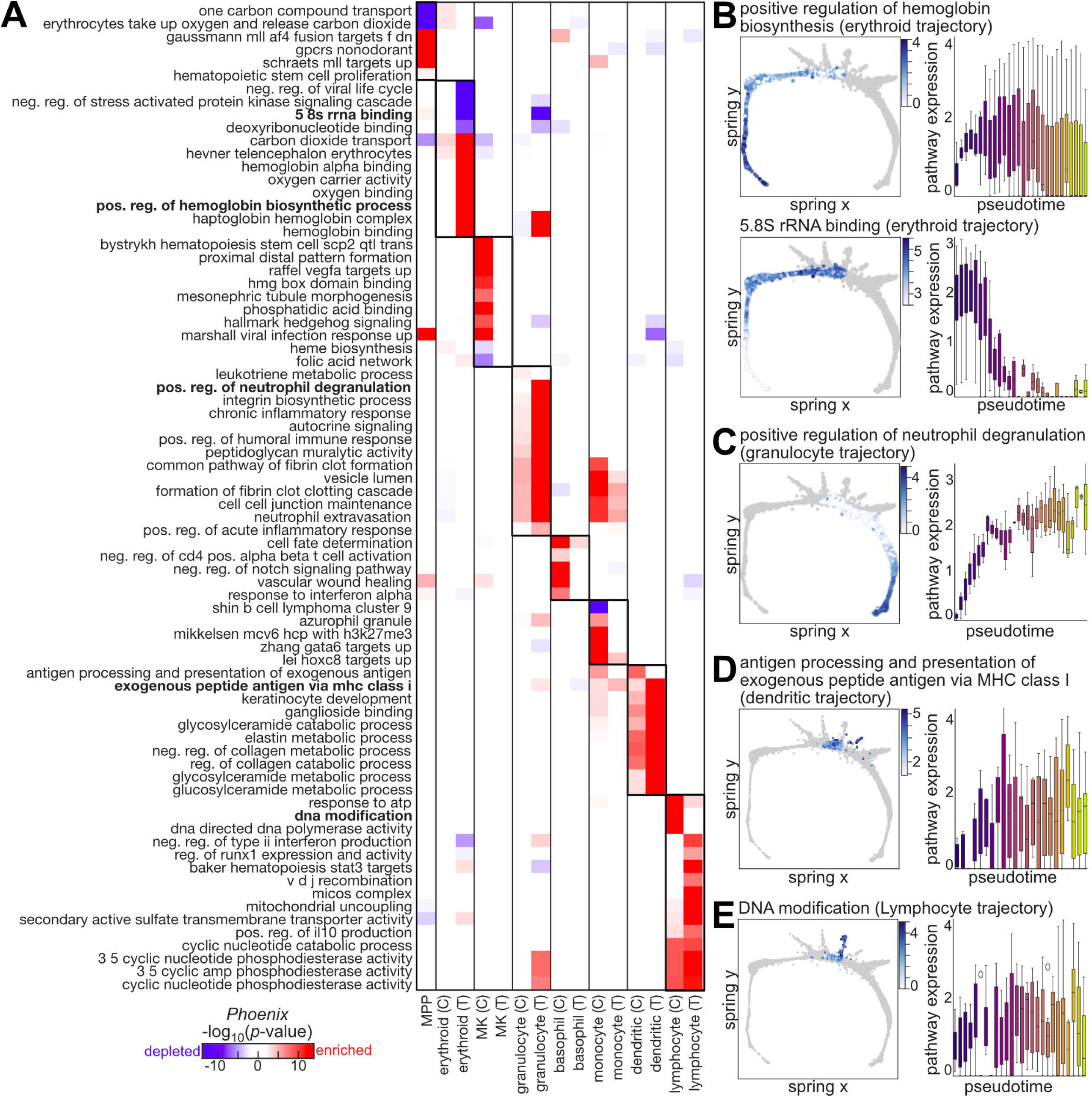
Pathways associated with mouse hematopoietic cell-types and trajectories. **(A)** Matrix of *p*-values calculated by *Phoenix* for the enrichment of top selected pathways (rows) within cell-types (noted by C) and trajectories (noted by T) of mouse hematopoietic scRNA-seq data (columns). Color-scale represents significance (-log_10_(*p*-value)) and effect: red positive values for enriched pathways (positive effect size) and blue negative values for depleted pathways (negative effect size). **(B-E)** Detailed expression data from selected pathways (as noted on top, and labeled in bold in panel (A)) that were associated with specific mouse hematopoietic differentiation trajectories by *Phoenix*. Left: scRNA-seq UMAP with trajectory cells colored by the average expression of genes within the selected pathway (blue color-scale, as indicated). Right: boxplot of average expression of genes within the selected pathway (y-axis) across pseudotime bins (x-axis). Box color represent pseudotime (purple: early; yellow: late). Central line represents the median, box edges are 25th and 75th percentiles, whiskers extend to largest/smallest value except outliers; outlier points are plotted individually.

Together, these results demonstrate *Phoenix*’s ability to identify significant and contextually relevant pathways in scRNA-seq data of hematopoietic differentiation across both cell types and trajectories.

### *Phoenix* expands biological associations compared to other tools

We compared *Phoenix* with several other pathway analysis tools (Mootha et al. 2003; Subramanian et al. 2005; Aibar et al. 2017; Franchini et al. 2023; Fan et al. 2016). While *GSEA* provides a class level analysis similar to *Phoenix*, in other tools (GSDensity, AUCell, Pagoda2, scGSEA) we aggregated per-cell scores into class-level results using *t*-test statistics to calculate a *p*-value, and average score difference to calculate an effect size. To perform a trajectory level comparisons, we calculated the correlation of per-cell scores with cells’ pseudotimes within each trajectory, using *t*-test statistics to calculate a *p*-value and correlation coefficient as the effect size (**Methods**, **Table S5**). In absence of external ground-truth pathway labels, we compared tools via global statistics and expected biological relationships.

The various tools identified different numbers of biologically relevant pathways in human and mouse hematopoietic differentiation (**Fig. 5A**). The number of predictions depended on size of the analyzed group of cells in some tools, but not in others (**Supplemental Fig. S6A**). In *Phoenix*, dependence only existed for trajectory analysis. Cell-type or trajectory specificity of predictions also varied between tools. *Phoenix* generally provides larger fractions of cell-type-specific predictions, but lower fractions of trajectory-specific associations (**Fig. 5A**). Overlap between predictions made by different tools was generally higher across cell-types than trajectories, but overall overlapping fractions were low (**Fig. 5B**). This limited agreement between all tools tested was consistent across individual cell types (**Fig. 5C**). Indeed, each tool provided a unique set of enrichments, and in some cases, one tool dominated enrichments identified for specific cell-types or trajectories. *Phoenix* and scGSEA dominated unique enrichments within trajectories. For example, scGSEA dominated enrichments within mouse and human MK and mast cell trajectories, while *Phoenix* dominated enrichments within mouse erythroid, monocyte and basophil trajectories, and human granulocytes and dendritic cells (**Fig. 5C**). Some of *Phoenix*’s unique associations provided additional biological insights. For example, in mouse it specifically linked basophils to dynamics of plasma lipoproteins, dendritic cells to adipocyte development, and correctly associated neutrophil-mediated immunity with granulocytes and monocytes. These are pathways that *GSEA* misassigned to erythroid and MK cells. *Phoenix* further suggested potentially novel regulatory roles, including regulation of various cells type differentiation and transdifferentiation by MK cells. These findings highlight *Phoenix*’s ability to capture both established and previously unrecognized pathway-level programs with high biological relevance.

**Figure 5.**
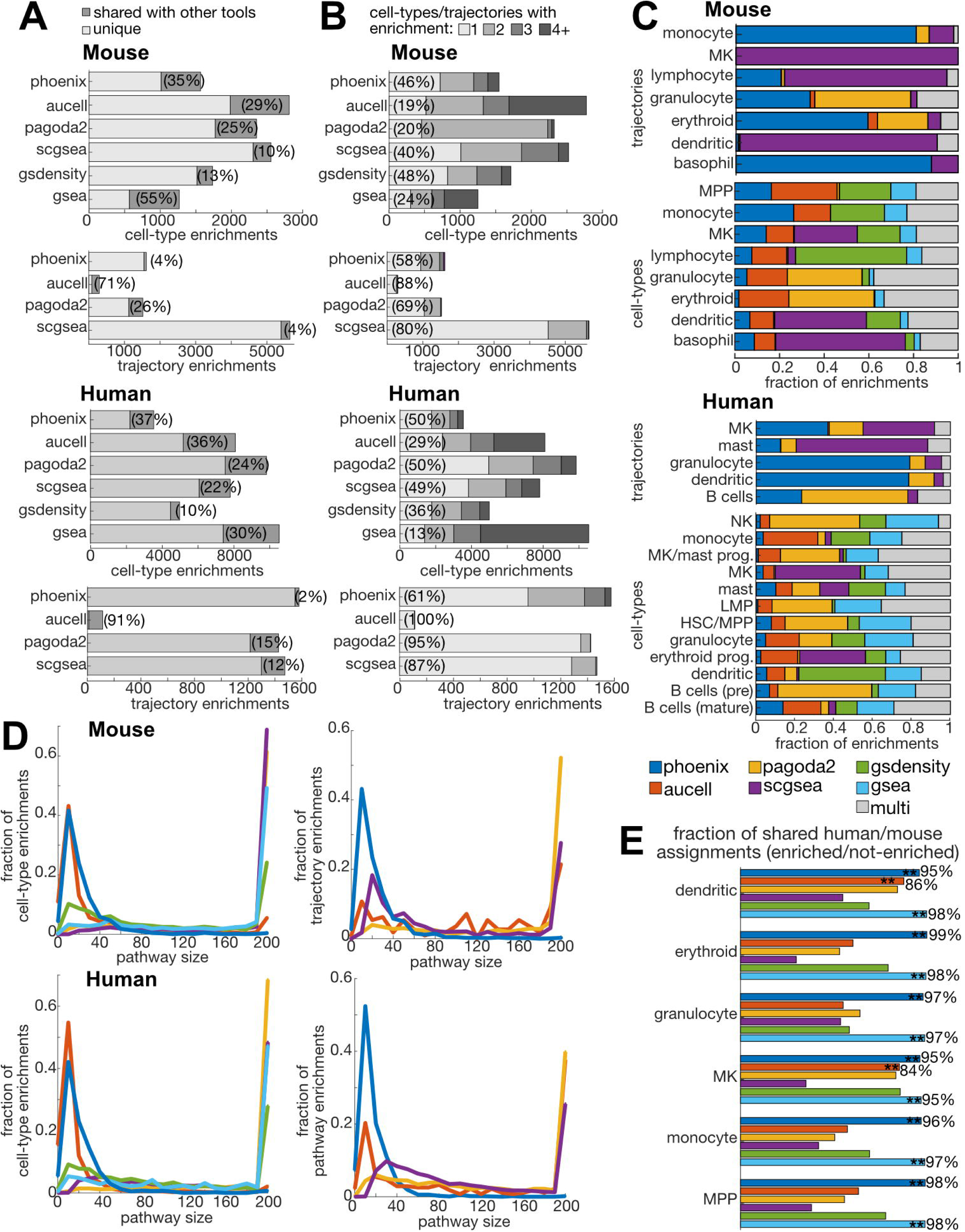
Analysis by *Phoenix* compared to other pathway analysis tools. **(A-B)** Number of cell-type (top) or trajectory (bottom) enrichments (x-axis) revealed by each of six pathway analysis tools for cell-types or four for trajectories (y-axis). Top: mouse, bottom: human. **(A)** Dark gray parts of bars represent the fraction out of annotations each tool revealed that are shared with at least one other tool. Total shared % is indicated in brackets. **(B)** Darker gray parts of bars represent which fraction of annotations each tool revealed are shared with one or more other cell-types/trajectories. Total unique % (not shared) is indicated in brackets. **(C)** Fraction of associations (x-axis) within cell-types (bottom) or trajectories (top) uniquely predicted by each pathway analysis tool (color coded, as indicated) or predicted by two or more of the tools (gray) in mouse (top) or human (bottom) hematopoiesis. **(D)** Distribution (y-axis, fraction) of sizes (number of genes assigned to pathway, x-axis) of enriched pathways identified by each pathway analysis tool (color coded, as indicated) in cell-types (left) or trajectories (right) in mouse (top) or human (bottom) hematopoiesis. **(E)** Percent of similarity in enrichments (x-axis) between mouse and human hematopoietic cell-types (y-axis) by six different pathway analysis tools (color coded, as indicated). Similarity is calculated by the number of similar assignments (enriched or not enriched) between human and mouse. As these could be affected by total number of assignments, random permutations of the assignments by each tool were performed to calculate a *p*-value for the specific association identified. Significant results (1% FDR corrected) are noted by **. Only *Phoenix* and GSEA were able to consistently reach significant similarities across all cell-types.

One of the main reasons for the low intersect with other tools is *Phoenix*’s unique ability to reveal enrichments within small pathways (**Fig. 5D**). Within cell-types AUCell was similarly linked to smaller pathways, but not within trajectories. *Phoenix*’s explicit use of a supervised ensemble model (random forest) can recover non-linear gene interactions within pathways that are not identified by other tools, and this ability could be particularly useful for small gene sets. Smaller gene sets likely contain fewer overlaps, and can offer higher resolution to detect specific biological activities within cells rather than more broad large gene sets. Indeed, pathway-level cross-species comparisons revealed *Phoenix*’s predictions were more consistent between human and mouse than those of most other tools (**Fig. 5E**), underscoring their biological robustness. Notably, GSEA reached similar consistencies as *Phoenix* in cross-species comparisons. These trends persist also when focusing only on top associations (top 10 associations within each cell-type/trajectory, **Supplemental Fig. S6B-C**) but overlaps between tools increase, as expected.

*Phoenix*’s ability to capture non-linear gene interactions is evident in pathways uniquely identified by it. For example, heme b biosynthesis was uniquely associated with mouse erythroid cells by *Phoenix*. SHAP (SHapley Additive exPlanations) analysis (Lundberg and Lee 2017) of the random forest model (**Fig. S7A**) revealed predominantly positive contributions from most pathway genes. However, the model also recovered a negative contribution to classification by *Alas1*, which has a minimal and largely redundant role in mouse erythroid cells. Contributions of the top three genes to this model were non-linear and depended on expression thresholds. *Phoenix* also uniquely associated the heme biosynthesis pathway with the mouse erythroid trajectory. The model recovered a clear temporal structure: high *Cpox* predicted early pseudotimes, whereas *Fech* levels were linearly associated with later pseudotimes (**Fig. S7B**). This aligns with known expression dynamics, where *Cpox* is active in erythroid progenitors (proerythroblasts), while *Fech* is induced later at the onset of hemoglobinization. In the same pathway the model also captured dependencies between genes, including for example, a reduced impact of *Fech* expression when *Cpox* levels were high (**Fig. S7B**). Finally, hematopoietic stem-cell proliferation was uniquely associated with mouse MPP progenitors by *Phoenix*. The learned model (**Fig. S7C**) revealed a strong positive effect to genes like *Cd34* (state marker) and *Pim1* (proliferation). Notably, while *Cd34* had an expression threshold contribution, *Pim1* expression contributed more linearly, and *Tsc22d1* had both a lower and upper threshold for high contribution.

Overall, our results demonstrate *Phoenix*’s ability to detect cell-type-specific, biologically meaningful pathways that other methods might miss within large single-cell datasets with multiple cell-types. The interpretable design of the *Phoenix* models further allows to extract meaningful biological insights regarding gene activity and interactions within pathways.

### *Phoenix* reveals lineage- and process-specific pathways in zebrafish embryogenesis

To further expose the broad utility of *Phoenix* for revealing contextually relevant biological pathways, we applied it to zebrafish embryogenesis, another system with complex, continuous cellular state transitions (**Supplemental Table S1**). Specifically, we analyzed zebrafish embryogenesis using a large scRNA-seq dataset (Farrell et al. 2018) comprising 29,830 cells organized into 25 differentiation trajectories (**Fig. 6A**), and evaluated pathway activities across 4,050 GO and KEGG terms.

**Figure 6.**
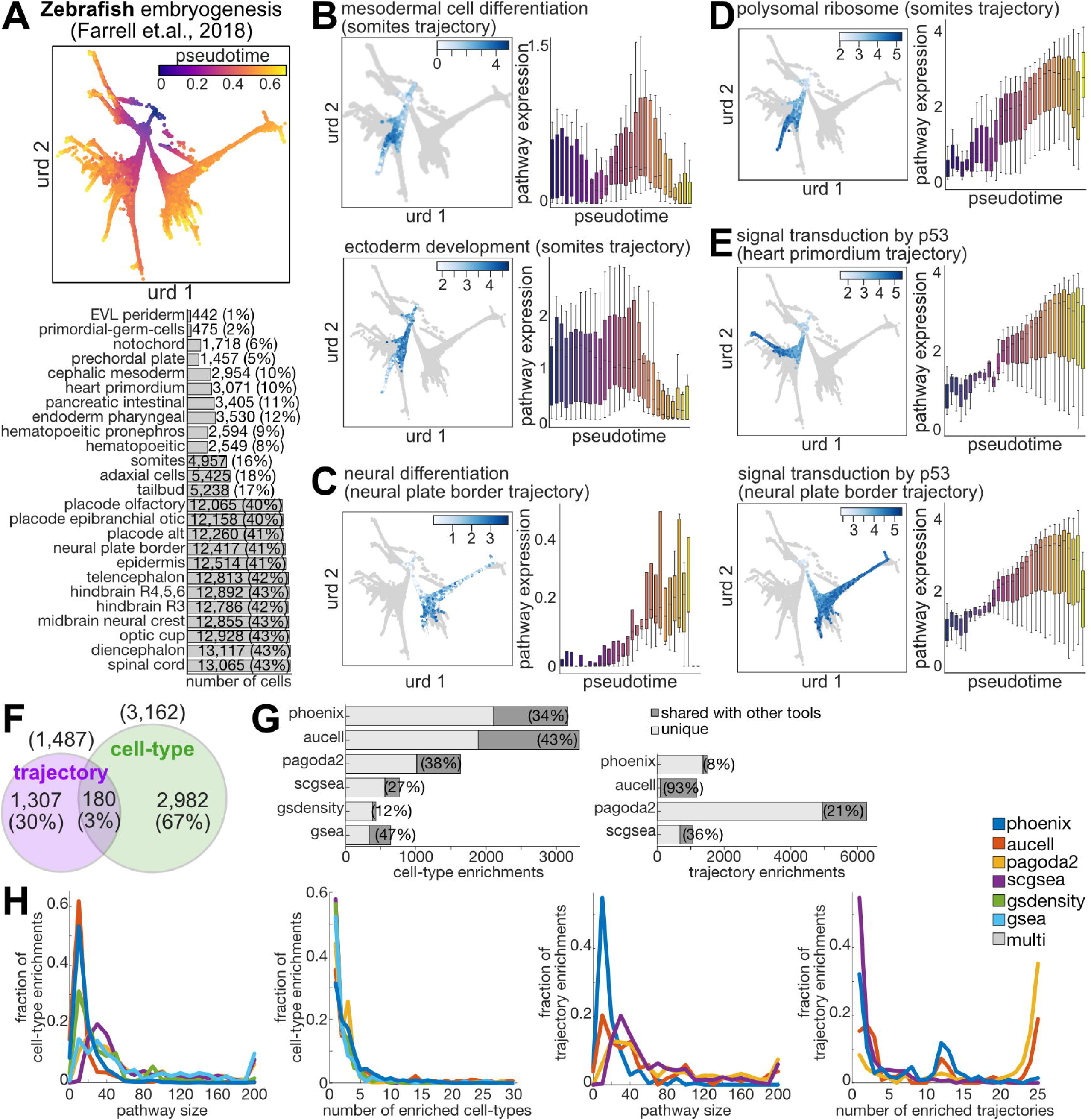
Pathways associated with cell-type specification during embryogenesis. **(A)** URD projection of zebrafish embryonic differentiation scRNA-seq data (Farrell et. al., 2018) with cells colored by pseudotime (purple: early, yellow: late). Number of cells assigned to each developmental trajectory is noted on bars. **(A-D)** Detailed expression data from selected pathways (as noted on top) that were associated with specific zebrafish embryonic development trajectories (Farrell et. al., 2018) by *Phoenix*. Left: scRNA-seq UMAP with trajectory cells colored by the average expression of genes within the selected pathway (blue color-scale, as indicated). Right: boxplot of average expression of genes within the selected pathway (y-axis) across pseudotime bins (x-axis). Box color represent pseudotime (purple: early; yellow: late). Central line represents the median, box edges are 25th and 75th percentiles, whiskers extend to largest/smallest value except outliers; outlier points are plotted individually. **(E)** Venn diagrams showing overlapping pathway associations by *Phoenix* with cell-types or matching trajectories in zebrafish embryonic development data. Total number of associations and percentages are indicated. **(F)** Number of cell-type (top) or trajectory (bottom) enrichments (x-axis) revealed by each of six pathway analysis tools for cell-types or four for trajectories (y-axis) in zebrafish embryonic development data. Dark gray part of bars represents the fraction out of annotations each tool revealed that are shared with at least one other tool. Total shared % is indicated in brackets. **(G)** Left: distribution (y-axis, fraction) of sizes (number of genes assigned to pathway, x-axis) of enriched pathways identified by each pathway analysis tool (color coded, as indicated) in cell-types (top) or trajectories (bottom). Right: distribution (y-axis, fraction out of total pathways identified) of the total number of cell-types (top) or trajectories (bottom) associated with each significant pathway identified (x-axis) by each pathway analysis tool (color coded, as indicated), as a measure of the uniqueness of associations.

Trajectory-level analysis identified 1,487 (1.5%) significant associations (FDR < 1%, effect size > 1.2; **Supplemental Table S6**), including 450 enriched (30%) and 1,037 depleted (70%) pathways, covering both housekeeping and developmental functions (**Supplemental Fig. S8**). Enrichments reflected expected lineage-specific programs, such as depletion of endodermal programs in ectodermal lineage cells, such as brain, optic cup and spinal chord, enrichment of mesodermal and depletion of ectodermal differentiation in somites (**Fig. 6B**), and enrichment of neural fate and neural differentiation in brain related lineages such as hindbrain, midbrain and neural plate border (**Fig. 6C**). Core cellular functions processes were also dynamically regulated across trajectories, including ribosome biogenesis and translation (**Fig. 6D**), p53 signaling, and DNA repair (**Fig. 6E**), consistent with previous reports of widespread remodeling of housekeeping programs during embryogenesis (Leesch et al. 2023).

Cell-type-level analysis, based on 41 cellular subpopulations defined by unique trajectory combinations (**Supplemental Table S7**), revealed 2,982 additional associations (**Fig. 6F**, **Supplemental Table S6**), encompassing both housekeeping and developmental functions (**Supplemental Fig. S9**). These included expected lineage programs: mesodermal fate enriched in mesodermal lineages (e.g., notochord, prechordal plate, axial mesoderm) and depleted in neural lineages; endodermal fate depleted in neural and other ectodermal lineages (e.g., epidermis); germ cells for germ plasm and P granules; and sensory placodes (otic/epibranchial, olfactory) and neural plate border enriched for peripheral nervous system differentiation and placode formation.

As with the hematopoietic analysis, in comparison to other tools *Phoenix* was able to identify trajectory-specific pathways of smaller size. While Pagoda2 revealed the largest number of trajectory associations (**Fig. 6G**), *Phoenix* and AUCell revealed the largest number of cell-type associations in this data, but remained tailored for cell-type specific pathways of smaller size (**Fig. 6H**, **Supplemental Table S5**).

Together, these results highlight *Phoenix*’s ability to uncover contextually relevant pathways in embryogenesis, offering high-resolution insights into both lineage- and process-specific pathways during continuous cellular state transitions.

## Discussion

In this study, we introduced *Phoenix*, a pathway analysis framework tailored for scRNA-seq data. *Phoenix* evaluates the relevance of biological pathways for both discrete cell-type distinctions and continuous cellular trajectories by training random forest models for classification or regression within a scRNA-seq dataset. These uncover biologically meaningful patterns that may be overlooked by other pathway analysis approaches.

Applying *Phoenix* to human and mouse hematopoietic differentiation data validated its capacity to recapitulate known biology and to discover new, contextually relevant associations. *Phoenix* identified pathways coherently reflecting the biological identities of specific cell types and trajectories. *Phoenix* identified both enriched or depleted pathways in discrete cell types and differentiation trajectories, highlighting multiple levels of cellular organization of biological processes. The zebrafish embryogenesis analysis further highlighted *Phoenix*’s versatility, and demonstrated its ability to reveal developmentally regulated pathways. Completeness and performance when analyzing data from non-model organisms will greatly depend on availability of high-confidence annotations.

Our comparative analysis against other commonly used tools further underscored *Phoenix*’s unique advantages compared to other methodologies. While all methods detected substantial numbers of pathways, exceeding in many cases the number of pathways reveal by *Phoenix*, overlap between them was low. *Phoenix* tended to identify smaller and more cell-type unique pathways, as its explicit use of machine-learning models enable it to focus on pathways that distinguish biological states, rather than those broadly active across cells. High consistency of *Phoenix* in pathway-level cross-species comparisons between human and mouse (notably distinct than gene-level orthology mapping) further underscored its biological robustness. In embryogenesis data, by complementing trajectory-based analysis with cell-type based analysis, *Phoenix* provided a more granular view, capturing developmental and housekeeping pathways relevant to dynamic cellular transitions, which were largely missed by other tools.

These results emphasize several strengths of *Phoenix*. First, its reliance on random forest models allows modeling of complex, non-linear gene interactions that traditional enrichment methods often miss, and discriminating between biological states even when expression changes are subtle. Interpretation technique of *Phoenix* models elicit internal relationships between pathways’ components and gene activities, further enhancing biological interpretability. Second, by explicitly integrating user-defined trajectories and cell-type annotations, *Phoenix* offers customized optimization for different datasets. Third, its dual focus on discrete and continuous cellular contexts aligns with the biological reality of single-cell data, where cells can both occupy distinct states and traverse dynamic processes. Because cellular processes often manifest as an interplay between discrete cell states and continuous progression, jointly modeling cell-type specificity and pseudotime dependence enables the identification of pathways whose roles could be missed or misinterpreted when each axis is considered in isolation, thereby improving both sensitivity and biological interpretability.

*Phoenix*’s limitations are rooted in its dependcy on the quality of trajectory and cell-type annotations, which themselves may be sensitive to data quality and preprocessing choices. Our current analysis primarily relied on the optimal trajectories inferred in the original studies, and robustness to alternative trajectory inference approaches still needs future evaluation. While effect size calculation aims to consider the entire trajectory, transient changes or complex expression dynamics along trajectories might remain obscure in this analysis. Additionally, while the random forest model captures complex interactions, it does not provide explicit mechanistic models of gene regulation; rather, it highlights statistically informative patterns that warrant further experimental validation.

Looking forward, *Phoenix* opens several avenues for advancing single-cell functional genomics. *Phoenix* is not limited to the analysis of discrete cell types, but can evaluate any predefined cellular grouping into cell populations within single-cell datasets, including for example, disease and control states, genetically or chemically perturbed subsets, and other experimental conditions. Similarly, regression-based analysis can integrate alternative continuous properties beyond pseudotime, such as spatial information to capture tissue context or spatial activity gradients. Large knowledgebases are also not the only source for pathway annotations. Approaches such as *Phoenix* can also bridge functional gene sets derived from perturbation-based assays, including CRISPR screens (Li et al. 2023; Xia et al. 2023) and MPRAs (Agarwal et al. 2025; Lagunas et al. 2023), to single-cell transcriptomic data by providing a principled framework for mapping experimentally defined regulatory elements or gene modules onto endogenous cellular states. *Phoenix*’s explicit use of a supervised ensemble model, interpretable design and focus on small concise gene sets can be particularly useful in these contexts, enabling detection of subtle or combinatorial effects, and eliciting internal relationships between pathways’ components. Its focus on compact gene sets improves robustness when transferring signatures across modalities, making it well suited for identifying the cellular contexts in which perturbation-derived signals are functionally active. By ranking informative genes within pathways, *Phoenix* offers hypotheses for key regulatory drivers that can be prioritized in downstream studies.

In summary, *Phoenix* provides a robust, flexible, and biologically insightful framework for pathway analysis in scRNA-seq data. Its capacity to reveal cell-type and trajectory-specific pathway enrichments, even in the presence of complex and subtle gene expression patterns, makes it a valuable addition to the single-cell analysis toolkit, facilitating deeper understanding of cellular heterogeneity and dynamic processes in health and disease.

## Methods

### *Phoenix*: pathway analysis of cell-types and developmental trajectories

*Phoenix* determines the contribution of functional pathways and other gene lists to the unique transcriptional profiles of cell-types or trajectories in scRNA-seq data. To do that, *Phoenix* trains a classification or a regression model using scaled scRNA-seq expression values from a specific group of genes (“pathway”) to classify cells in the experiment into cell-types (classification) or to predict their pseudotime along a trajectory (regression). *Phoenix* uses a Random Forest (RF) model, a robust learning method that generates an ensemble of decision trees, where each tree is trained on a random subset of the input data. As an ensemble approach, it reduces risk of overfitting and improves generalization, making it effective for complex datasets like scRNA-seq. *Phoenix* assesses the models’ performance by comparing it to that of other gene sets of the same size. *Phoenix* is implemented in an open-source Python package, and all computations are based on *scikit-learn* functions (Pedregosa et al. 2011).

#### Normalization

*Phoenix* applies *StandardScaler* to normalize provided single-cell gene expression levels (*Z*-score normalization).

#### Cell-types classification models

*Phoenix* treats cell-type analysis as a classification task, and approaches it in two ways: (1) multi-class classification, where all cell-types are simultaneously predicted, and (2) binary classification, where each cell-type is classified separately to determine whether a cell belongs to that specific type or not. The target variable is a cell-type label per cell, encoded into numerical categories by *LabelEncoder*. *Phoenix* trains a RF classifier (*RandomForestClassifier* with the ‘entropy’ criterion and default parameters) with 20 decision trees of a maximum depth of 20 each, for both multiclass and binary classification. The model is trained with 10-fold cross-validation, and the median of the 10 *cross_val_score* values is used as the final training score.

To account for potential imbalances in the number of cells of each cell-type, *Phoenix* uses a weighted F1-score with normalized inverse class frequencies (each class score is weighed by the inverse of its frequency), thus ensuring that the model’s performance on minority classes is appropriately considered and not overshadowed by majority classes. Specifically:

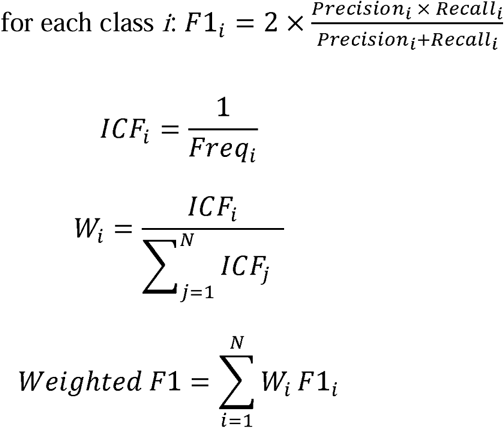

#### Pseudotime regression models

*Phoenix* treats trajectory analysis as a regression between gene expression values within each cell and pseudotimes assigned to cells along a trajectory. The target variable is a scaled pseudotime per cell, transformed per trajectory into values between 0 and 1 by *MinMaxScaler*. This transformation allows to compare the results between different trajectories and datasets. *Phoenix* trains a RF regression model (*RandomForestRegressor* with the ‘squared_error’ criterion and default parameters) with 20 decision trees of a maximum depth of 20 each. The model is trained with 10-fold cross-validation, and the median of the 10 *cross_val_score* values is used as the final training score. To evaluate regression *Phoenix* uses the negative Root Mean Squared Error (RMSE) score. Whereas MSE is more sensitive to outliers or significant deviations since it assigns more weight to larger errors by squaring the residuals before averaging, RMSE (the square root of MSE) returns the error measure to the same scale as the original data, providing a more intuitive interpretation. In our case, where the target pseudotime values are scaled between 0 and 1, RMSE is especially useful as it directly corresponds to the data’s scale. Specifically:

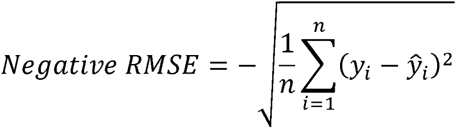

#### Significance evaluation

For each dataset, *Phoenix* generates a background distribution of classification and regression prediction scores, and fits these scores a Gamma distribution parametric model. *Phoenix* assigns a *p*-value for each pathway as the probability of the pathway score based on the CDF of the Gamma distribution fitted to a gene set with a size most closely matching the pathway size. It applies a Benjamini-Hochberg multiple hypothesis correction across all tested pathways (across all cell-types and all trajectories). *Phoenix* provides two alternatives for Gamma distribution evaluation.

##### (1) Random mode

In this mode *Phoenix* selects 150 random gene sets of a specified size from the data, and apply the above modeling procedures to obtain 150 prediction scores. Gene set sizes used are 2, 5, 10, 15, 20, 25, 30, 40, 60, 80, 100, 150, 200 (total = 13 sizes, 150 scores for each). An Interquartile Range (IQR) based filtering is applied to exclude extreme values that could distort the distribution. IRQ is defined as the difference between the 25th percentile (*Q1*) and the 75th percentile (*Q3*) of the data, and values below a lower bound of *Q1 - 1.5*×*IQR* or above an upper bound of *Q3 + 1.5*×*IQR* are considered outliers. Finally, a Gamma distribution (shape, location, and scale parameters) is fitted to filtered background scores (using the *stats.gamma* module from *SciPy* (Virtanen et al. 2020)).

Our analysis (**Supplemental Fig. S1**) showed that the Gamma distribution fits the background *Phoenix* prediction scores by evaluating the goodness of the Gamma fit with a Kolmogorov-Smirnov (KS) test, confirming that as the sample size increased, the fit to the distribution improved. The fit stabilized around sample sizes of 150-200, suggesting that increasing the sample size beyond this range does not significantly improve the fit.

##### (2) Real mode

In this mode *Phoenix* uses the pathway scores of the entire pathway set to estimate background Gamma distributions, and can be used if number of pathways is 2,000 or more. Pathway scores from the data are divided into groups of 150 pathways based on their size. These scores are analyzed as described above to estimate a Gamma distribution (shape, location, and scale parameters), but using a more stringent outlier filtering with k=0.5 (instead of 1.5 in random mode), defining a lower bound of *Q1 - 0.5*×*IQR* and an upper bound of *Q3 + 0.5*×*IQR*. The resulting Gamma distributions are used to estimate the significance of pathways in the group.

Our analysis (**Supplemental Fig. S2**) showed that the Gamma distribution fits using real pathway scores provide similar results as with random scores for most cell-types and trajectories, with significantly reduced run times in this mode.

#### Effect size

For each pathway enrichment, *Phoenix* also calculates an effect size which is defined by the fold-change in mean expression (geometric mean with natural log) of pathway genes. For cell-type enrichments, fold-change is defined between the tested cell-type and all other cells in the dataset. For trajectory enrichments, fold-change is defined between the 20% of cells with the earliest pseudotime, and temporal groups of 20% of cells along the trajectory using a sliding window approach, ending with the latest 20% of cells. We exclude from this calculation redundant windows with a mean pseudotime that is very close (< 0.05) to an earlier window along the trajectory. *Phoenix* reports the largest effect size and time selected by this calculation. Effect sizes are subsequently standardized by mean (for each cell-type or trajectory).

#### Feature selection

*Phoenix* offers an option to perform feature selection on each pathway prior to model fitting, to select a subset of pathway genes that are most relevant for predicting cell-types or pseudotimes. In this procedure, *Phoenix* defines a size (*N_G_*) per pathway by selecting the closest size to the desired *fraction* of genes per pathway parameter (e.g., 0.75) from the list of background distribution sizes (as defined by each background mode). In Real mode, the median size within each group is used. It uses *SelectKBest* to select the *N_G_* highest scoring genes (after normalization) within each pathway, using the ANOVA F-statistic calculated by *f_regression* for pseudotimes and *f_classif* for cell-types for selection. For the analysis performed in this work, feature selection was not employed (keeping 100% of pathway genes).

#### Per-gene contributions

For each pathway enrichment, *Phoenix* reports per-gene contribution diagnostics based on the random-forest model learned (using *feature_importances_* attribute of the RF model), to reveal which genes contributed most significantly to the classification or regression task. Per-gene importance is summarized as the median feature importance across folds, and genes are ranked in descending order of this median importance. By default, *Phoenix* filters out pathways in which >50% of pathway genes contributed below a 5% low-importance threshold.

### Single-cell RNA-seq datasets

We analyzed 3 single-cell RNA-seq (scRNA-seq) datasets (**Supplemental Table S1**): hematopoietic stem and progenitor cell differentiation in human (Ranzoni et al. 2021) and mouse (Tusi et al. 2018), and early embryonic development of zebrafish (Farrell et al. 2018). Published single-cell data, dimensionality reduction coordinates and cell-type and trajectory annotations were used when available, or used *Slingshot* (Street et al. 2018) to define lineages and pseudotimes.

#### Human hematopoiesis

Count matrix and cell-type annotations were obtained from https://www.ebi.ac.uk/gxa/sc/experiments/E-MTAB-9067. We retained only cells sourced from ‘bone marrow’ that passed quality control, and excluded any ‘cycling’ cells or ones with unspecified annotations. We combined granulocyte progenitor and granulocyte 1, 2 and 3 into a single ‘granulocyte’ category, and monocyte 1 and 2 into a ‘monocyte’ class. Also, we combined the pre-B and B progenitor cells, annotated as ‘pro-B’. Similar to the original paper, we excluded endothelial cells (31 single cells) from the cell-type classification analysis. Gene symbols were obtained by converting Ensembl identifiers. *Slingshot* (Street et al. 2018) was used to define lineages, utilizing the first 10 principal components.

#### Mouse hematopoiesis

Wild-type count matrices, metadata, and cell annotations were downloaded from https://kleintools.hms.harvard.edu/paper_websites/tusi_et_al (the ‘Kit+ Bone marrow, basal’ link). Cell-type annotations were derived from probability scores in the metadata and published marker genes, with cells filtered based on quality criteria. Given that the published pseudotime values ranged from zero (representing the earliest cell) to -300 (representing the most differentiated cell), we applied min-max normalization and reversed the scale so positive values represent developmental progress. This scaling adjustment does not impact the analysis since pseudotime is scaled during preprocessing and their numerical values are not inherently meaningful. The provided dimensionality reduction coordinates from *SPRING* (Weinreb et al. 2018) were used for visualization.

#### Zebrafish development

An *URD* object (Farrell et al. 2018) containing raw counts, cell identities, pseudotime values and developmental tree coordinates was retrieved from https://singlecell.broadinstitute.org/single_cell/study/SCP162/single-cell-reconstruction-of-developmental-trajectories-during-zebrafish-embryogenesis. As the original paper mostly focused on trajectory analysis, we used this information to generate cell-type classifications. We grouped cells that shared identical trajectory assignments, meaning that each category included all cells belonging to a specific subset of trajectories while excluding all others. This resulted in 41 distinct cell-types (**Supplemental Table S5**), which were subsequently used by *Phoenix* for cell-type analysis.

### Single-cell RNA-seq data processing

#### Gene and cell filtering

Single-cell RNA-seq data preprocessing was performed to ensure high-quality input for downstream analysis using *SCANPY* (Wolf et al. 2018). We excluded low-quality cells with fewer than 100 expressed genes, and lowly expressed genes with fewer than 500 unique molecular identifiers (UMIs) and expressed in less than 5 cells. Data were normalized (using *SCANPY noralize_total* with target_sum=1e^4^) to account for variations in sequencing depth across cells, followed by a logarithmic transformation to stabilize variance. Finally, to focus the analysis on the most informative features, we selected the top 5,000 genes based on their mean expression levels.

#### Dimensionality reduction

To visualize the annotated single-cell data in 2D, we either used provided dimensionality reduction coordinates (mouse and zebrafish data), or reduced dimensionality using Uniform Manifold Approximation and Projection (UMAP) with default parameters (human).

### Pathway annotations

We define a ‘pathway’ as a set of genes that share a functional annotation. Pathways used in this study were obtained from MSigDB (Liberzon et al. 2011; Subramanian et al. 2005), KEGG (Kanehisa et al. 2025; Kanehisa and Goto 2000) and Gene Ontology (The Gene Ontology Consortium et al. 2000, 2023). We utilized the Python libraries *GSEApy* (Fang et al. 2023) and *BioServices* (Cokelaer et al. 2013) for queries. For human and mouse datasets, we used MSigDB annotations, which include also KEGG and GO annotations. Since MSigDB does not support zebrafish, only KEGG and GO were queried for zebrafish. We filtered each pathway to include only genes observed in the filtered expression data, and retained only pathways with at least 2 genes. To identify matching genes between datasets and pathways, we considered all three possible cases – lowercase, uppercase, and capitalized (first letter uppercase, rest lowercase).

### Defining marker gene-sets

For human (Ranzoni et al. 2021) and mouse (Tusi et al. 2018) hematopoiesis datasets, we use a list of marker genes per cell-type as defined in the reference papers.

### Reporting *Phoenix* results

*Phoenix* was applied to each scRNA-seq dataset using the following parameters background_mode=random; repeats=150; set_fraction=1.0; min_set_size=2; fdr_threshold=0.01; corrected_effect_size_threshold=1.2; importance_lower_threshold=0.05; importance_gene_fraction_threshold=0.5; processes=60. Resulting *p*-values represent 1% FDR corrected values, across all pathways and cell-types of a specific dataset combined. Results were filtered to include only pathways with effect size >= 1.2, and in which >50% of pathway genes contributed above a 5% low-importance threshold.

### Comparison to other pathway analysis tools

We compared *Phoenix* to several publicly available pathway analysis tools.

#### (1) GSEA

*GSEA* (Mootha et al. 2003; Subramanian et al. 2005) is one of the most popular tools to determine the relevance of biological pathways within gene expression data. It uses a statistical test to estimate the enrichment of pathway genes within the top or bottom of an ordered list of genes. To apply *GSEA* to single-cell data, we ordered genes by the fold-changes (FC) in their cell-type specific expression, and applied *GSEA* to calculate significance of pathway annotations (using FGSEA v1.24.0 in R (Korotkevich et al. 2021)). FC was calculated by the ratio of average expression levels in one cell-type against those in other cell-types (plus a small constant of 10^−4^ added to avoid division by zero). *P*-values were adjusted by FDR. *GSEA* was run using its R implementation. Human pathways were queried via the *msigdbr* package (v10.0.2). For mouse and fish, because *msigdbr* uses ortholog mapping for non-human species, in order to ensure consistency with *Phoenix*, we used the exact annotation lists generated by *Phoenix* (via *gseapy* or *bioservices*). We selected pathways that showed significant results (p<0.01) in at least one cell-type. Since *GSEA* is primarily designed for discrete classifications, we did not analyze trajectories.

#### (2) GSDensity

*GSDensity* (Liang et al. 2023) is a method for detecting coordinated functional gene sets, specifically designed for scRNA-seq data, and does not require prior cell-type partitioning. GSDensity was cloned from https://github.com/KChen-lab/gsdensity and applied using a Seurat (Satija et al. 2015) object constructed with minimal filtering (min.cells=3, min.features=200). We applied *GSDensity* to calculate binarized Pathway Activity Level (PAL) for each pathway in each cell in the data, splitting the cells into two groups with high (“positive”) and low (“negative”) relevance to the pathway. The analyses for human, mouse, and fish all used the annotation lists from *Phoenix*. We used a hypergeometric *p*-value and FDR correction to test the frequency of “positive” cells within each cell type compared to cells not assigned to it. We selected pathways that showed significant results (p<0.01) in at least one cell-type, and did not analyze trajectories.

#### (3) scGSEA

*scGSEA* (Franchini et al. 2023) extends Gene Set Enrichment Analysis to single-cell RNA-seq data by computing enrichment scores at the individual cell level rather than across bulk samples. It ranks genes within each cell and evaluates whether genes from a given pathway are coordinately up- or downregulated within that cell using a running-sum enrichment statistic similar to *GSEA*. We applied scGSEA using gficf (v2.0.0) with nmf.k = 50, after filtering out small gene sets (<10 genes). We compared the per-cell scores between cells assigned to each cell-type to those not assigned to it. Since this approach produced many significant results, we used *t*-test statistics and multiple hypothesis correction with the Bonferroni method, to stringently focus only on the most significant pathways, and selected pathways that showed significant results (p<0.01) in at least one cell-type. In addition, we calculated the difference between score averages within a cell-type and cells not assigned to it as a measure of “effect size”, and apply a filter to include only results within the top 30-th percentile of their distribution. For trajectory analysis, we calculated the correlation of per-cell scores with cells’ pseudotimes within each trajectory. We calculated significance by *t*-test statistics and Bonferroni correction, and selected pathways that showed significant results (p<0.01) in at least one cell-type. We retained only results that also had a correlation coefficient above 0.7.

#### (4) Pagoda2

*Pagoda2* (v1.0.13) is a pathway and gene set overdispersion analysis framework, which expands the original PAGODA algorithm (Fan et al. 2016). It identifies pathways that show significant variability (overdispersion) across cells beyond technical noise expectations. Aggregation of scores per cell-type or trajectory was performed as described above.

#### (5) AUCell

*AUCell* (Aibar et al. 2017) evaluates pathway activity at the single-cell level by calculating the Area Under the Curve (AUC) for the recovery of pathway genes within the ranked expression profile of each cell. For each cell, genes are ranked by expression, and *AUCell* measures whether genes from a given set are enriched at the top of this ranking, producing a continuous activity score per pathway per cell. We applied AUCell (v1.20.2) after filtering low-count cells (<300 counts) and lowly expressed genes (<10 cells), followed by CPM normalization and log1p transformation. Aggregation of scores per cell-type or trajectory was performed as described above.

To minimize differences in the number of tested pathways across tools, which arised from the way each tool obtains and filters pathway data, pathways in tested tools were matched to *Phoenix* lists whenever needed. Notably, pathways in *Phoenix* were also filtered to exclude those without any genes in the filtered expression data (after retaining the top 5,000 genes), resulting in the removal of roughly 30–51% of pathways, while other tools have different filtering steps, resulting in fewer pathways removed in pre-filtering.

### Comparison between species

Pathway analysis was applied independently to each species, using its annotated pathways. Pathways were compared between species based on their annotated names and ids. Cell-type classes were defined to refer to a similar cell-population between species by comparing their pre-defined identities within each species.

### SHAP analysis

SHAP (SHapley Additive exPlanations) analysis is a method to explain how predictions are made by machine learning models that can be applied to any machine learning model as a post hoc interpretation technique (PonceLBobadilla et al. 2024). SHAP analysis uses game-theoretic Shapley values by considering the potential input variables (genes in a pathway) as “players” in a collaborative game: working together to determine each individual predicted value (classification into cell-types or regression into pseudotime), and quantifies how important each input variable is to a model for making predictions (Lundberg and Lee 2017). SHAP analysis calculates contributions for each of the model’s input variables, and their sum, plus a fixed base value, reproduces the machine learning model’s predictions for each instance. We applied SHAP analysis to random forest models trained using the same configuration as in *Phoenix*, and computed SHAP values using *shap* (v0.51.0) by applying *TreeExplainer*. Our analysis was performed both on the full dataset, and by training on 80% of the input data (cells) and computing SHAP values on the 20% held-out cells. Results were visualized using *shap*’s beeswarm and dependence plots.

## Code availability

*Phoenix* is implemented in Python (version 3.11) and can be accessed at https://github.com/NachmaniLab/Phoenix. We used Python 3.11 with the following libraries: *NumPy* (1.26.4), *Pandas* (2.2.2), *Dask* (2024.7.1), *Scikit-learn* (1.5.1), *SciPy* (1.14.0), *seaborn* (0.13.2), *Matplotlib* (3.9.1), *SCANPY* (1.10.2), *GSEApy* (1.1.3), and *BioServices* (1.11.2). To ensure reproducibility, we managed the software environment using a virtual environment (Conda or Python venv) and fixed random seeds across all runs. The *Phoenix* repository also provides documentation on how the analyses in this paper are run, and additional custom scripts and code required to reproduce the work are provided as **Supplemental Code**.

## Competing interest statement

The authors declare that they have no competing interests.

## Acknowledgements

We thank Yotam Drier for helpful discussions and critical reading of the manuscript. This research was supported by the European Research Council Horizon 2020 (grant 852451 to MR, and grant 101075607 to DN). MR and DN conceived and designed the project. YH wrote the *Phoenix* code. MR and YH performed the downstream analyses. All authors participated in writing the manuscript.

## Notes

### Competing Interest Statement

The authors have declared no competing interest.

### Summary of Updates

Algorithm adjusted and analysis revised accordingly.

