## Supplemental Figures for "A unified analysis of cell-type and trajectory-associated pathways in single-cell data using Phoenix"

#### Supplemental Fig. S1

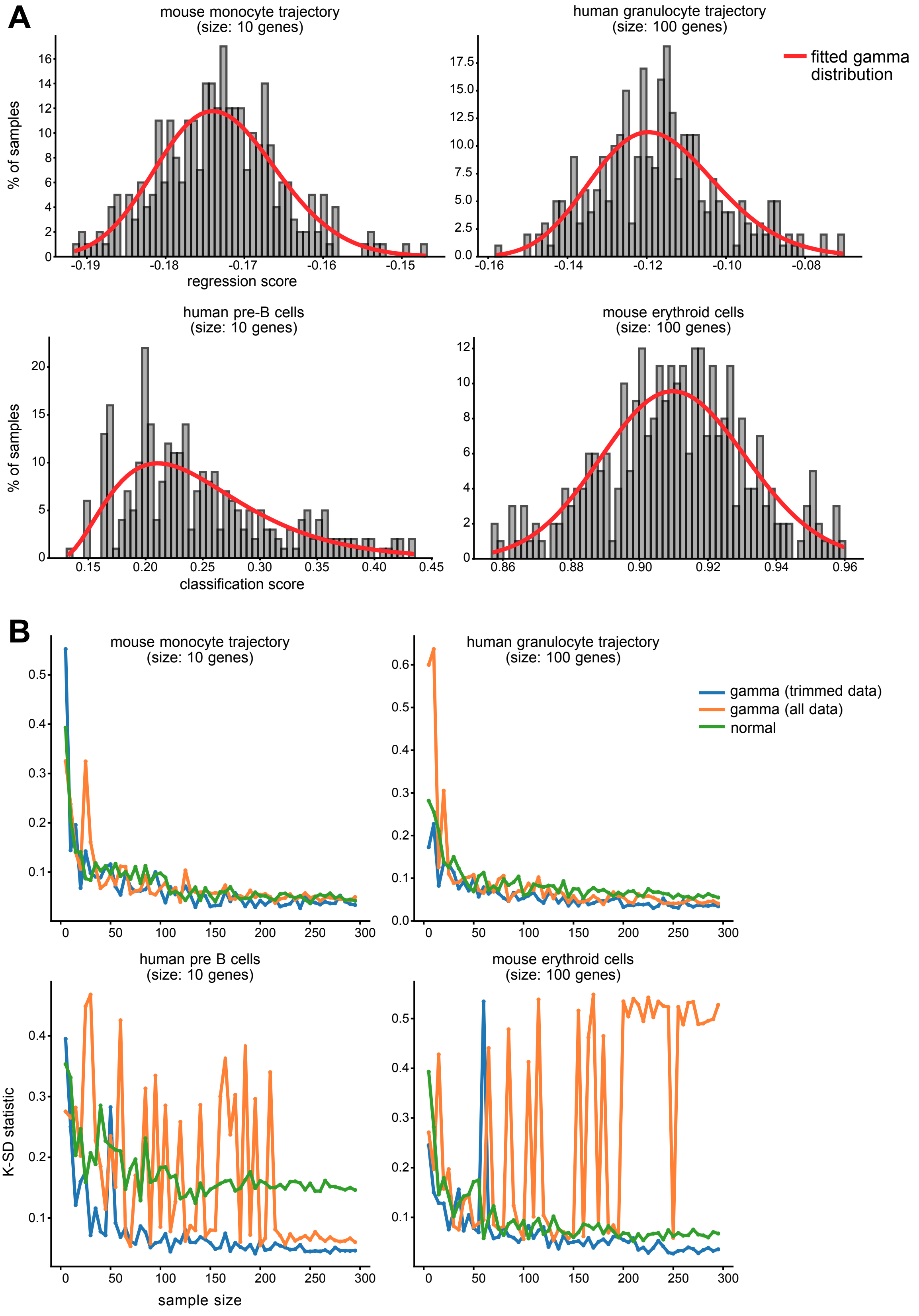

Supplemental Fig. S1. Gamma distribution fit for background prediction scores.

**(A)** Histograms of classification and regression scores using random gene sets across different datasets, gene set sizes, cell-types, and trajectories, overlaid with fitted Gamma distributions (after IQR-based filtering). **(B)** Kolmogorov–Smirnov (KS) statistics comparing the fit of Gamma distributions (with and without IQR filtering) and normal distributions across increasing sample sizes. Gamma fits on trimmed data outperform both unfiltered Gamma and normal fits, and stabilize around 150–200 samples.

### Supplemental Fig. S2

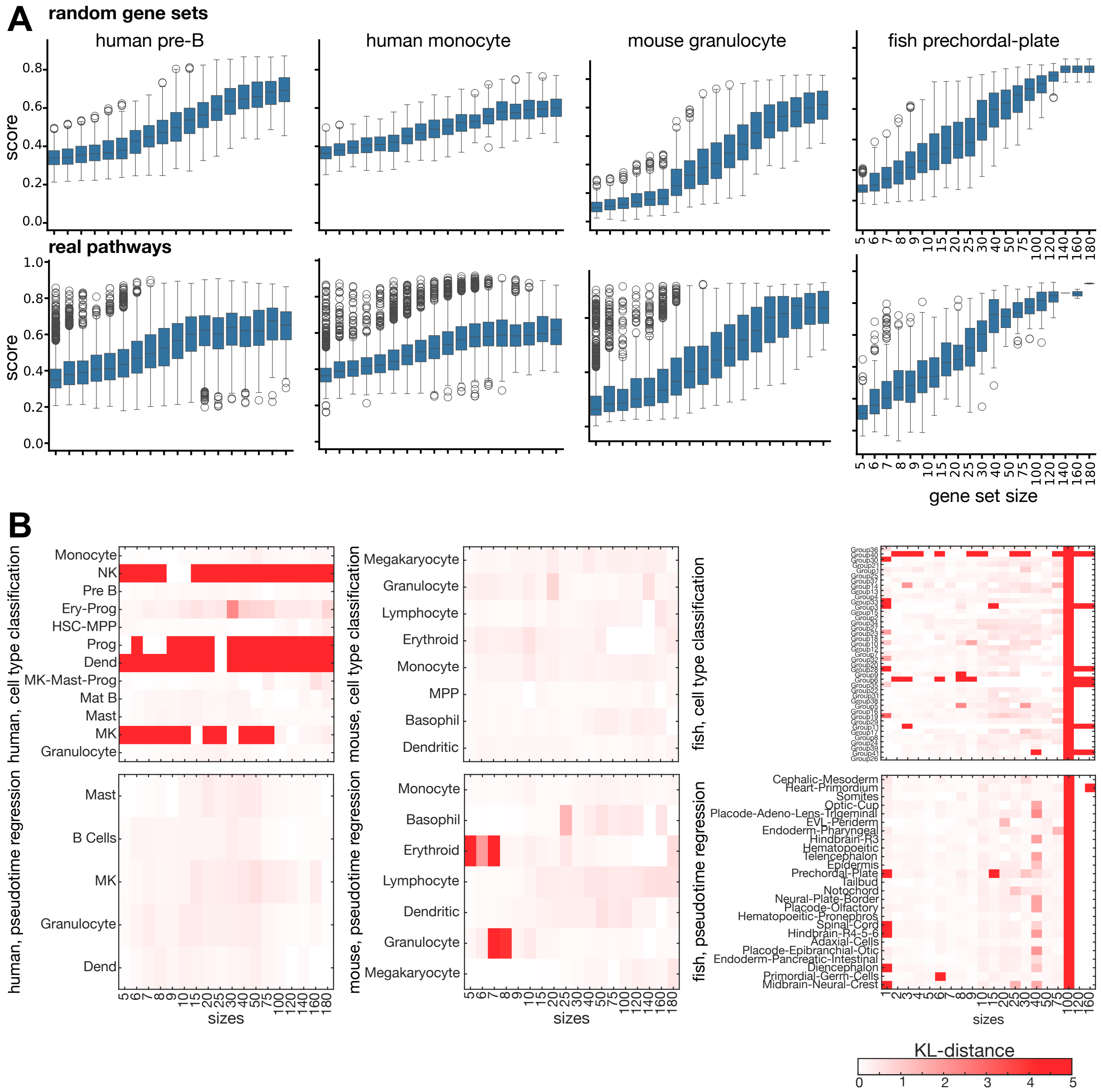

Supplemental Fig. S2. Score calculations and evaluation of gamma distributions by random gene sets compared to real pathways.

**(A)** Score distributions for random gene sets (left) and real pathways of similar sizes (right). These values are used for estimating a gamma distribution. **(B)** KL-distance (red intensity) between gamma distributions estimated by random gene sets and real pathways of similar sizes. Distances remain low in most cases, except few specific cell-types.

### Supplemental Fig. S3

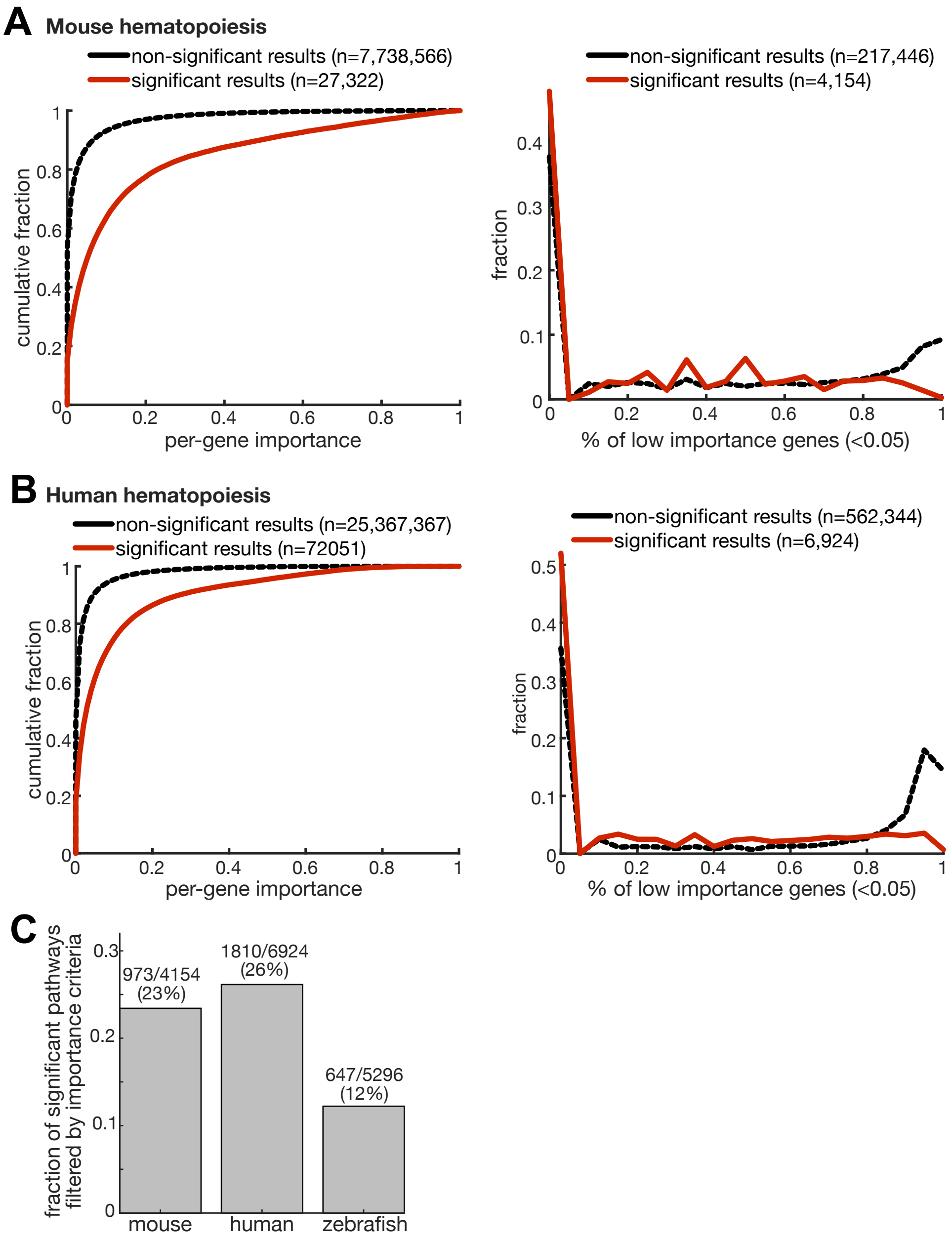

Supplemental Fig. S3. Analysis of gene importances in random-forest models

**(A-B)** Left: cumulative distribution (y-axis) of per-gene importance scores (x-axis) based on random forest models learned by *Phoenix*, in significant (red) and non-significant associations across all cell-types and trajectories. Right: distribution (y-axis) of % of low-importance genes (x-axis) across all tested associations, defined as importance of less than 5% in random forest models learned by *Phoenix*, in significant (red) and non-significant associations; **(A)** in mouse, **(B)** in human hematopoietic datasets. **(C)** Fraction of associations (y-axis) with a significant *p*-value that were filtered due to a high fraction (50% or more) of low-importance genes in each of three tested datasets (x-axis). Numbers filtered and percentages are noted on top.

### Supplemental Fig. S4

#### A Runtime benchmark on a small dataset

| | total runtime | % time reduced (vs. original) | pathway scoring (% out of run) | background calculation (% out of run) | wall-clock time | dataset |
|---|---|---|---|---|---|---|
| random gene set based background | 8h 0m 23s | 43% | 52% | 47% | 2h 26m 1s (5 processes) | 1,708 cells 1,855 pathways 2 cell-types 1 trajectory |
| pathway-based background | 4h 48m 50s | 66% | 98% | 0% | 1h 6m 0s (5 processes) | |

#### B Runtime and memory benchmark on full-scale single-cell datasets

| background calculation | total memory | total runtime | wall-clock time (60 processes) | dataset |
|---|---|---|---|---|
| random gene sets | 5.11 GB | 91h 25m 6s | 4h 19m 9s | **mouse**: 13,850 pathways 9 cell-types with 4,751 cells 7 trajectories with 5,705 cells |
| pathway-based | 5.11 GB | 73h 40m 34s | 3h 25m 56s | |
| random gene sets | 3.62 GB | 417h 57m 34s | 12h 32m 45s | **human**: 31,626 pathways 13 cell-types with 2,939 cells, 5 trajectories with 6,767 cells |
| pathway-based | 3.61 GB | 356h 1m 22s | 11h 15m 59s | |
| random gene sets | 21.86 GB | 838h 2m 21s | 47h 22m 59s | **fish**: 2,792 pathways 42 cell-types with 30,163 cells 25 trajectories with 189,685 cells |
| pathway-based | 21.77 GB | 237h 56m 21s | 6h 13m 23s | |
| random gene sets | 11.71 GB | 214h 57m 25s | 10h 36m 19s | **fish, reduced size**: 2,792 pathways 42 cell-types with 15,968 cells 25 trajectories with 22,960 cells |
| pathway-based | 11.70 GB | 78h 33m 40s | 3h 0m 38s | |

* number of cells across trajectories is larger due to assignment of cells to multiple trajectories

Supplemental Fig. S4. Benchmarking of runtime and memory requirements in *Phoenix*.

**(A)** List of runtime benchmark results on a small dataset, consisting of one trajectory (with 1,708 cells) and 2 cell-type (with 1,549 and 159 cells), and selected a subset set of 1,855 pathways of variable sizes for testing. Benchmarking using this set confirmed that the overall runtime was reduced by 66% when using pathways themselves as background. Moreover, dividing the run into 5 parallel processes on a computational cluster shortened the overall runtime to 2.5 wall-clock hours with background calculation and 1 hour without it. **(B)** Evaluation of memory and runtimes on the 3 full datasets used in this study, and a subset of the full zebrafish dataset, using 60 parallel processes on a computational cluster. Runs using either random gene set or pathway based (faster) background calculation.

### Supplemental Fig. S6

**A**

correlation between number of enrichments and number of cells in **cell-type**

| Mouse | Human |
|---|---|
| phoenix (0.04) | phoenix (0.05) |
| aucell (**0.96**) | aucell (**0.65**) |
| pagoda2 (**0.99**) | pagoda2 (-0.23) |
| scgsea (-0.40) | scgsea (-0.33) |
| gsdensity (-0.50) | gsdensity (0.30) |
| gsea (**0.97**) | gsea (0.11) |

correlation between number of enrichments and number of cells in **trajectory**

| Mouse | Human |
|---|---|
| phoenix (**0.93**) | phoenix (-0.06) |
| aucell (**0.77**) | aucell (-0.29) |
| pagoda2 (**0.65**) | pagoda2 (-0.56) |
| scgsea (-0.36) | scgsea (0.02) |

**B** Mouse

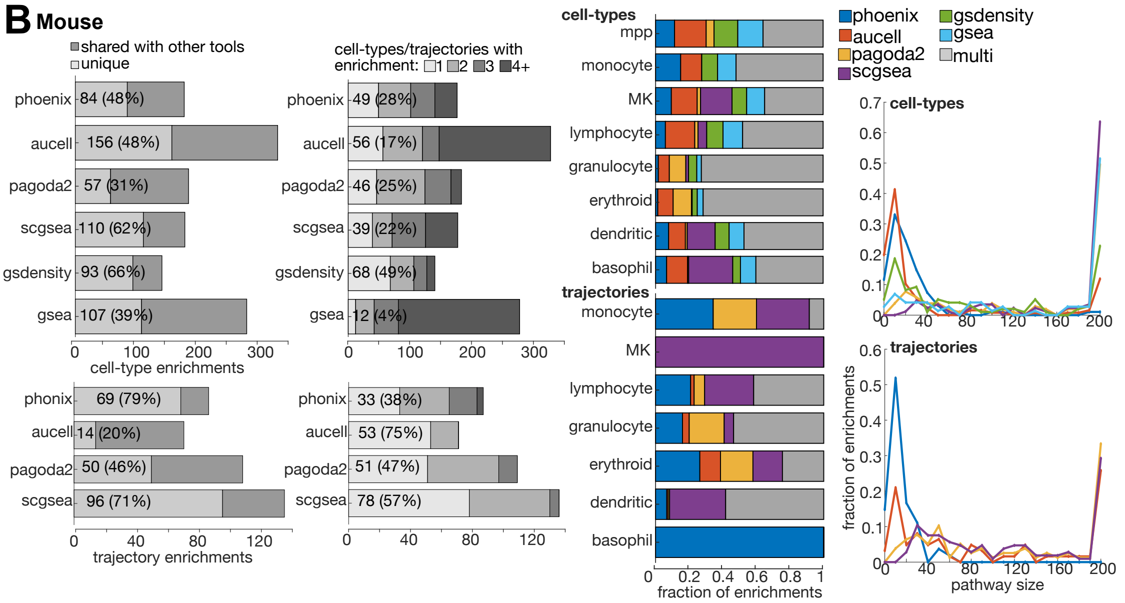


**C** Human

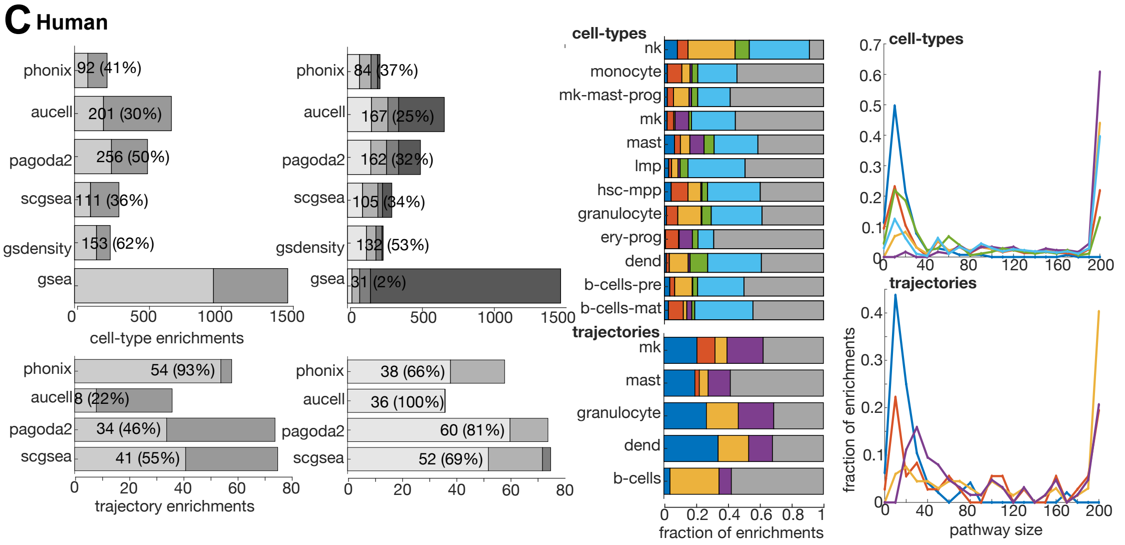

Supplemental Fig. S5. *Phoenix* analysis of hematopoietic differentiation.

**(A-B)** Matrix of *p*-values calculated by *Phoenix* for marker gene sets (columns) within cell-types (top) and trajectories (bottom) of hematopoietic scRNA-seq data (rows) reveals their association with expected cell-types within scRNA-seq data. First row of the matrix represents the classification of all cell-types in the data (multi-class classification). Color-scale represents significance ($-\log_{10}$(*p*-value)) and effect: red positive values for enriched pathways (positive effect size) and blue negative values for depleted pathways (negative effect size). **(A)** Human data (Ranzoni et. al., 2021) analyzed by human (left) or mouse (right) marker sets. **(B)** Mouse data (Tusi etl al., 2018) analyzed by mouse (left) or human (right) marker sets. **(C-D)** Number of enriched (red) or depleted (blue) pathways in human **(C)** or mouse **(D)** cell-types and trajectories.

### Supplemental Fig. S5

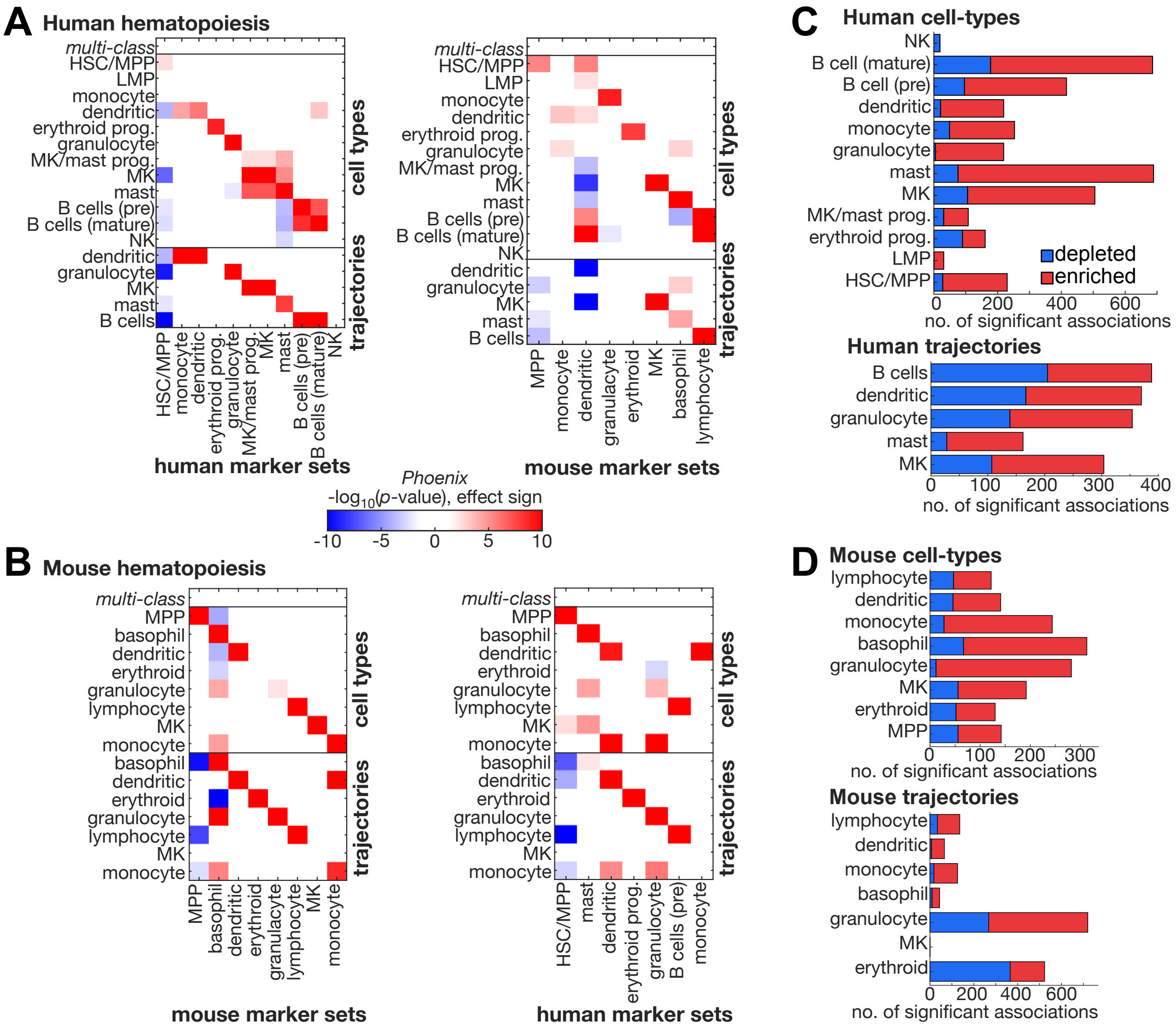

Supplemental Fig. S6. Analysis by *Phoenix* compared to other pathway analysis tools.

**(A)** Calculated *Pearson* correlation coefficients between number of associations identified by each pathway analysis tool in each cell-type (left) or trajectory (right) and the size (number of cells) of each group. High correlation coefficients could indicate a stronger dependency of number of associations identified with size of group. **(B-C)** Comparison between different pathway analysis tools, using the top 10 annotations (by *p*-value) for each cell-type or trajectory in **(A)** mouse or **(B)** human. Left: number of cell-type (top) or trajectory (bottom) enrichments (x-axis) revealed by each of six pathway analysis tools for cell-types or four for trajectories (y-axis). Dark gray part of bars represents the fraction out of annotations each tool revealed that are shared with at least one other tool. Total shared % is indicated in brackets. Middle-left: Darker gray parts of bars represent which fraction of annotations each tool revealed are shared with one or more other cell-types/trajectories. Total unique % (not shared) is indicated in brackets. Middle-right: fraction of associations (x-axis) within cell-types (bottom) or trajectories (top) uniquely predicted by each pathway analysis tool (color coded, as indicated) or predicted by two or more of the tools (gray). Right: distribution (y-axis, fraction) of sizes (number of genes assigned to pathway, x-axis) of enriched pathways identified by each pathway analysis tool (color coded, as indicated) in cell-types (top) or trajectories (bottom).

### Supplemental Fig. S7

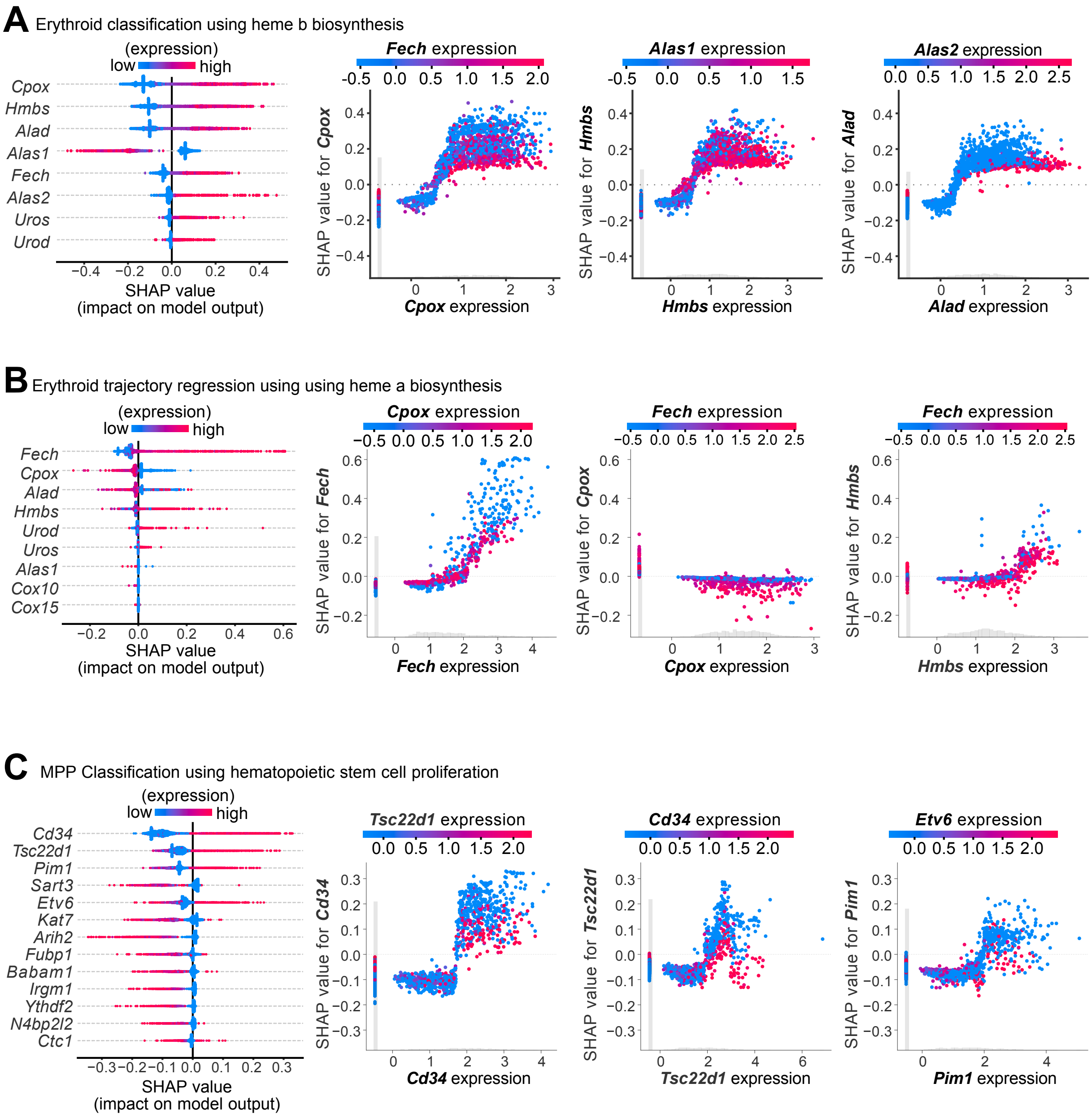

Supplemental Fig. S7. SHAP analysis of random forest models learned by *Phoenix*.

SHAP analysis results for the following models, representing associations uniquely identified by *Phoenix*: **(A)** classification model for mouse erythroid cells using heme b biosynthesis pathway; **(B)** regression model for mouse erythroid trajectory using heme a biosynthesis pathway; **(C)** classification model for mouse MPP cells using hematopoietic stem cell proliferation. Left: Beeswarm plot showing SHAP values (x-axis) for each gene (y-axis). Each dot represents a single-cell in the data, and is colored by the expression value of the gene within that cell (red: high expression, blue: low expression). Right: dependence plot for top 3 genes, showing the relationship between a gene's expression values (x-axis) and the model's predicted outcomes, represented by SHAP values (y-axis). Each dot represents a single-cell in the data, and is colored by the expression value of a strongly interacting second gene in the cell, as indicated on top (red: high expression, blue: low expression). The insert histogram above the x-axis (gray) represents the distribution of the gene’s raw expression values.

### Supplemental Fig. S8

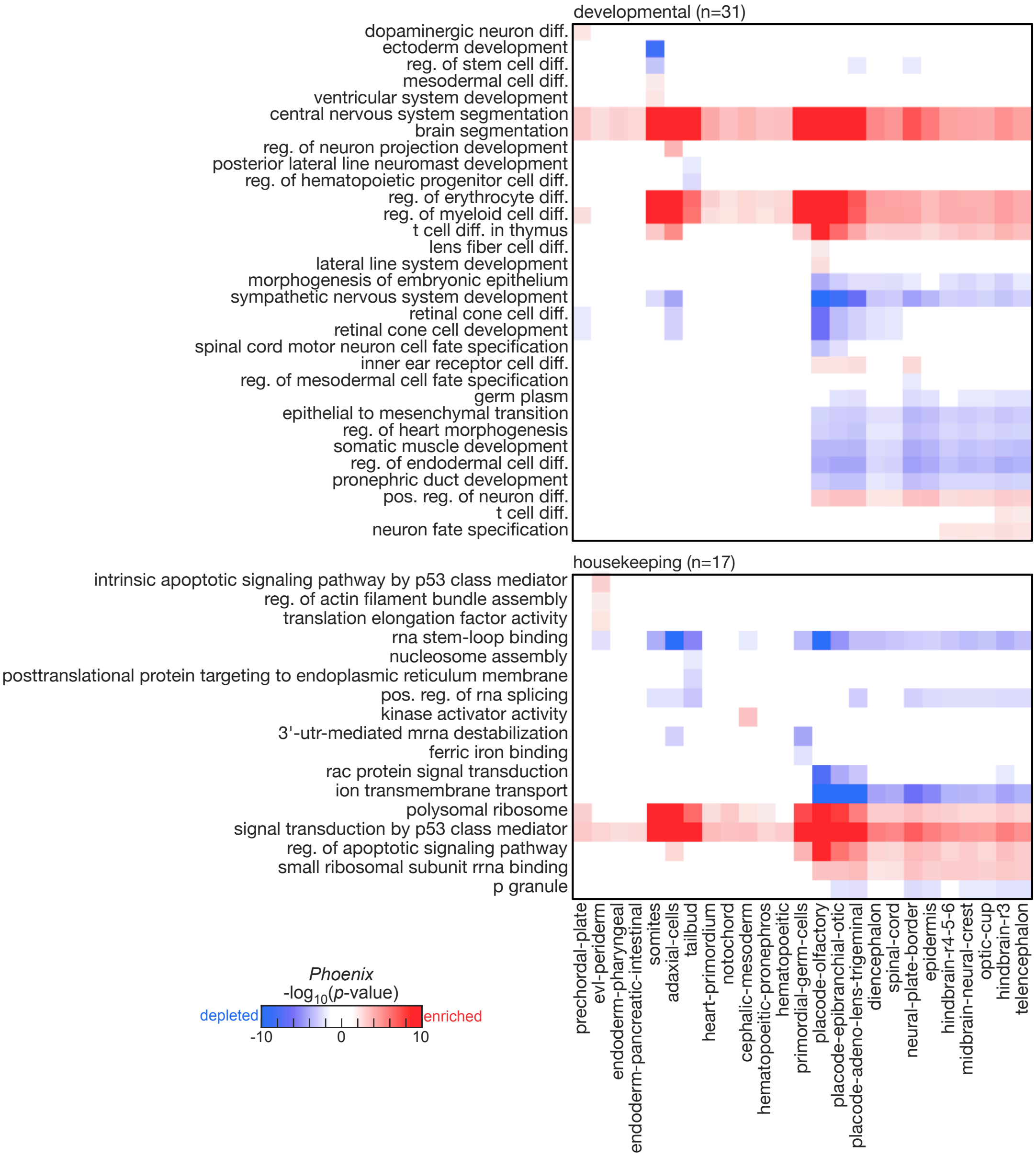

Supplemental Fig. S8. Pathways associated with developmental trajectories of zebrafish embryogenesis.

Matrix of *p*-values calculated by *Phoenix* pipeline for the enrichment of selected pathways (rows) within zebrafish embryonic developmental trajectories (columns). Pathways are divided by developmental (top) and housekeeping (bottom) functions. Color-scale represents significance (-$\log_{10}$(*p*-value)) and effect: red positive values for enriched pathways (positive effect size) and blue negative values for depleted pathways (negative effect size).

### Supplemental Fig. S9

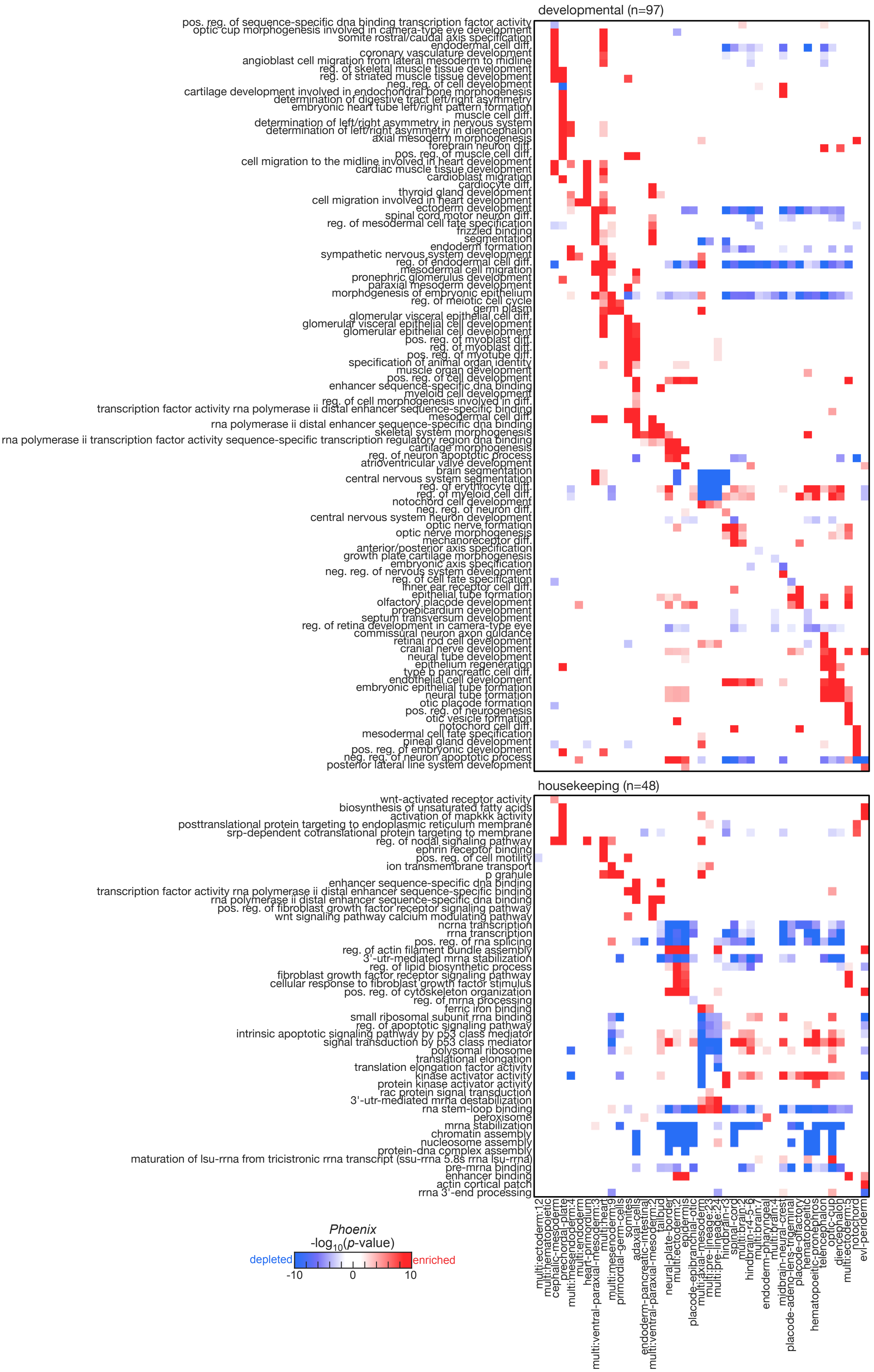

Supplemental Fig. S9. Pathways associated with developmental cell-types of zebrafish embryogenesis.

Matrix of *p*-values calculated by *Phoenix* for the enrichment of selected pathways (rows) within zebrafish embryonic developmental cell-types (columns). Pathways are divided by developmental (top) and housekeeping (bottom) functions. Color-scale represents significance (-$\log_{10}$(*p*-value)) and effect: red positive values for enriched pathways (positive effect size) and blue negative values for depleted pathways (negative effect size).

#### Supplemental Analysis

##### Runtime and memory requirements

Empirical p-value estimation used in *Phoenix* leads to increased computation times. In its original version, for each new dataset *Phoenix* calculated background distributions using random gene sets of different sizes (200 scores x 25 different sizes, per cell-type or trajectory), and then fitted a gamma distribution to the resulting scores (per size).

In the revised manuscripts, we offer several important adjustments to reduce needed calculations and improve runtime. First, we reduced the number of set sizes used for score calculations from 25 to 13 different sizes. Analysis of estimated gamma distributions for 25 different sizes in our original implementation showed very similar results for similar sizes (**Fig. R1**), allowing us to reduce number of sizes without significantly affecting the results. If tested pathway set contains fewer sizes, *Phoenix* will only calculate the relevant subset of sizes.

Second, for runs with large pathway sets (~1,500 pathways or more), *Phoenix* now offers the option to rely on the pathway set itself to calculate background distribution, thus completely avoiding random gene set calculations. When pathway sets are large enough, we expect the bulk of pathways are not biologically meaningful, providing a background distribution for p-value estimation (with outlier filtering). For this calculation, *Phoenix* divides all input pathways into 13 bins by their size, and fit a gamma distribution per bin, using a more stringent outlier filtering. Analysis of fitted gamma distributions based on this approach confirmed similar models as random gene sets in most cases (**Supplemental Fig. S2**). This approach both shortens runtime by avoiding random gene-set calculations, and relies on real biological pathways for background calculation, offering possible enhanced resolution.

Our repository also provides detailed instructions and scripts for running the tool within a cluster environment, as a practical usage approach which is common practice for large datasets and significantly shortens runtimes.

These changes affected the final pathways reported, but overall ~70% of significant pathways were retained with the updated random gene-set background compared to our original results, and ~72% of significant pathways retained with the updated pathway-based background compared to random gene-set results (**Fig. R2**).

To improve memory requirements, we allow single-cell data to also be provided in a standard 10x Genomics MTX sparse format, which is commonly used for single-cell expression matrices and results in significantly smaller file sizes compared to a standard CSV format.

To evaluate the improvement of running time by these modifications, we constructed a small dataset, consisting of one trajectory (with 1,708 cells) and 2 cell-type (with 1,549 and 159 cells), and selected a subset set of 1,855 pathways of variable sizes for testing (**Fig. R3**, **Supplemental Fig. S4A**). Benchmarking using this set confirmed that 77% of runtime in the original implementation was dedicated to background calculation. The overall runtime was reduced by 43% in the new implementation (with fewer background sizes), and by 66% when using pathways themselves as background. Moreover, dividing the run into 5 parallel processes on a computational cluster shortened overall runtime to 2.5 wall-clock hours with background calculation and 1 hour without it.

Evaluation of memory and runtimes on the full datasets (**Supplemental Fig. S4B**) confirmed feasible runtimes for full-scale single-cell datasets, particularly using parallel processes on a cluster environment (notably, up to 10% differences in runtime can arise due to variation in availability of cluster resources).

**Transient pathway regulation along pseudotime**

We adjusted *Phoenix*'s effect size calculation in trajectory analysis. Rather than calculating only the difference in expression of pathway genes between 20% earliest and 20% latest cells in a trajectory, *Phoenix* also compares expression of the earliest cells to additional groups of cells along the trajectory using a sliding window approach, and reports the largest effect size and time selected by this calculation. This approach both captures transient changes and provides additional information on the time window with maximal change. While this change had minimal effect on calculation for most genes, it provided enhanced resolution in some cases (**Fig. R4**), shedding more light at the ability of *Phoenix* to address such cases.

### Figure R1

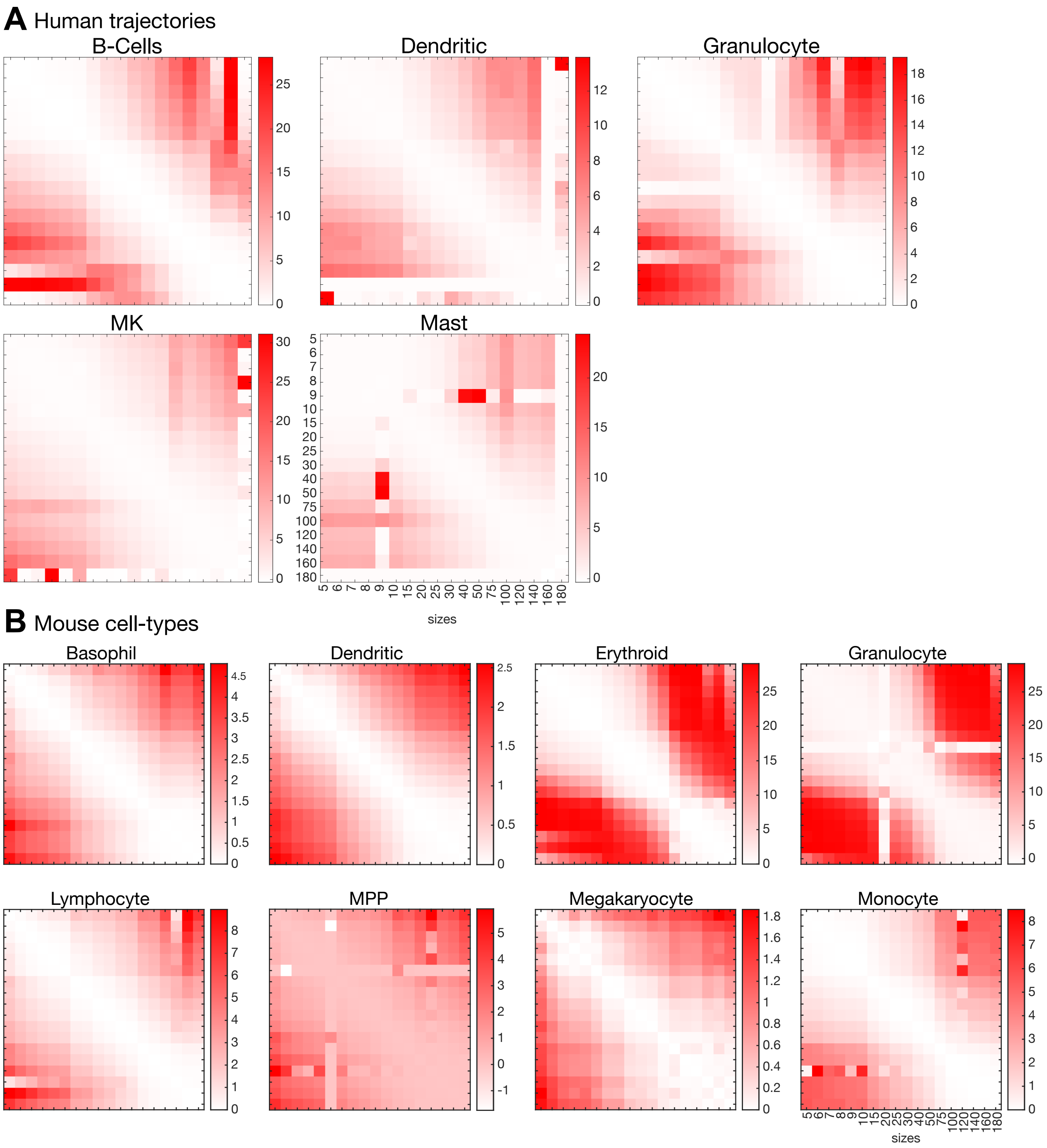


**Figure R1**. **Number of sizes required for background score calculations**. KL-distance (red intensity) between gamma distributions estimated by random gene sets of 25 different sizes. Distances between similar sizes (across diagonal) show very similar results, allowing us to reduce number of sizes without significantly affecting the results.

### Figure R2

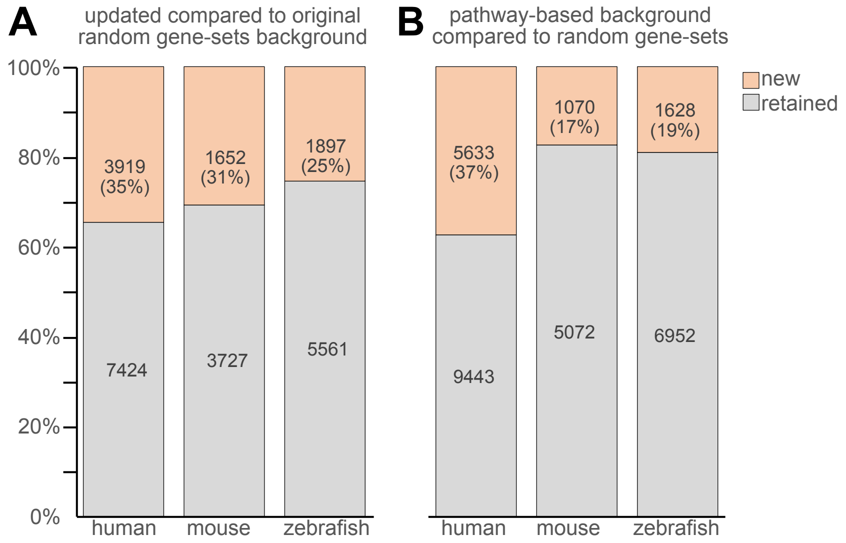


**Figure R2**. **Changes in pathway assignment in undated version of *Phoenix***. Bar plots showing % of significant pathways that were retained after changes introduced into *Phoenix* background calculation. **(A)** compared to original version (random background). **(B)** Compared to pathway-based background calculation.

### Figure R3

**Runtime benchmark on a small dataset**

| | total runtime | % time reduced (vs. original) | pathway scoring (% out of run) | background calculation (% out of run) | wall-clock time | dataset |
|---|---|---|---|---|---|---|
| original implementation | 14h 2m 39s | 0% | 22% | 77% | 2h 56m 41s (15 processes) | 1,708 cells<br>1,855 pathways<br>2 cell-types<br>1 trajectory |
| random gene set based background | 8h 0m 23s | 43% | 52% | 47% | 2h 26m 1s (5 processes) | |
| pathway-based background | 4h 48m 50s | 66% | 98% | 0% | 1h 6m 0s (5 processes) | |

**Figure R3. Benchmarking of runtime and memory requirements in *Phoenix*.** Table shows runtime benchmark results on a small dataset, consisting of one trajectory (with 1,708 cells) and 2 cell-type (with 1,549 and 159 cells), and selected a subset set of 1,855 pathways of variable sizes for testing. Benchmarking using this set confirmed that 77% of runtime in the original implementation was dedicated to background calculation. The overall runtime was reduced by 43% in the new implementation (with fewer background sizes), and by 66% when using pathways themselves as background. Moreover, dividing the run into 5 parallel processes on a computational cluster shortened the overall runtime to 2.5 wall-clock hours with background calculation and 1 hour without it.

### Figure R4

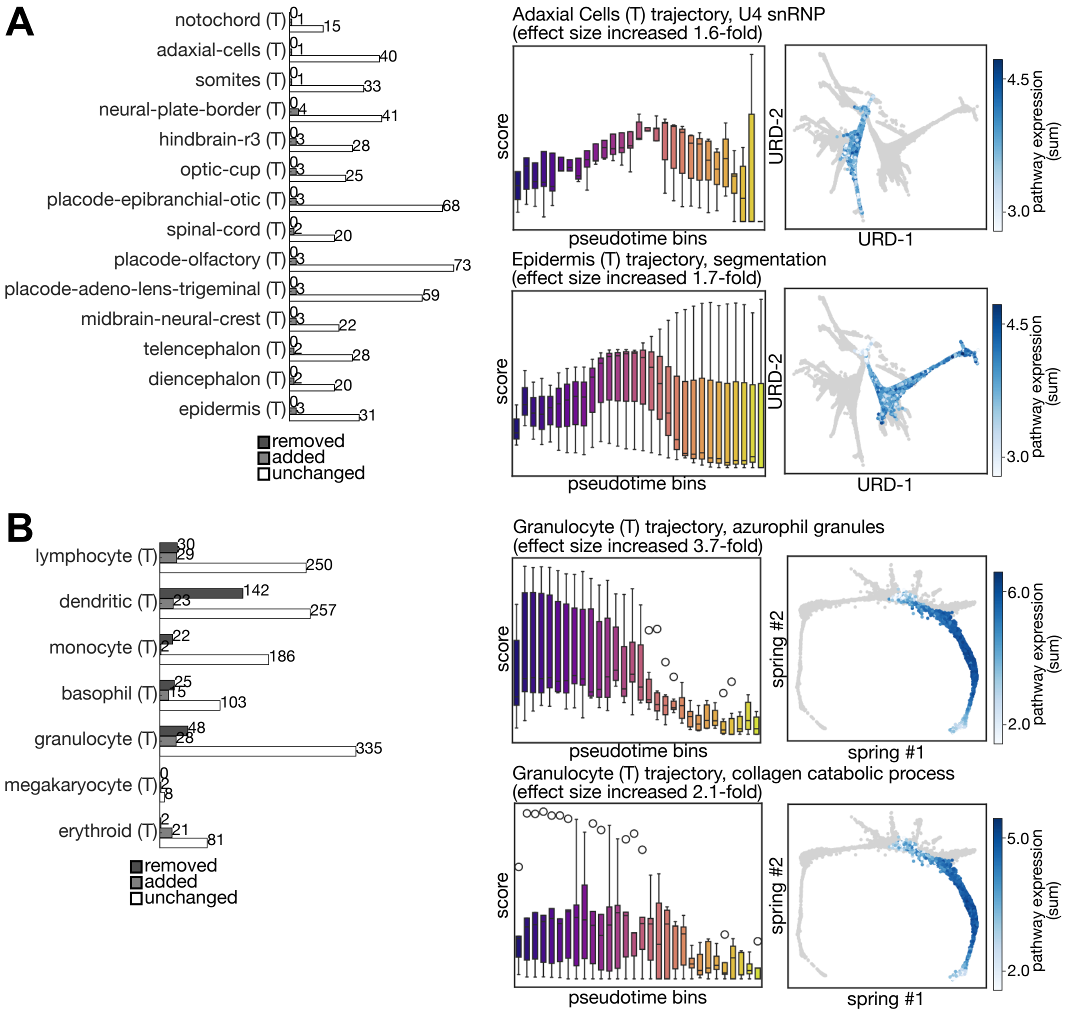


**Figure R4. Effect size calculation in pathways with transient changes in expression in trajectories**. Number of pathways which remain, removed or added as associated with each trajectory based on new effect size calculation compared to previous calculation. Most pathways do not change. **(A)** Zebrafish development data. **(B)** Mouse hematopoiesis data.
